# Elucidation of an anaerobic pathway for metabolism of L-carnitine-derived γ-butyrobetaine to trimethylamine in human gut bacteria

**DOI:** 10.1101/2021.01.25.428109

**Authors:** Lauren J. Rajakovich, Beverly Fu, Maud Bollenbach, Emily P. Balskus

## Abstract

Trimethylamine (TMA) is an important gut microbial metabolite strongly associated with human disease. There are prominent gaps in our understanding of how TMA is produced from the essential dietary nutrient L-carnitine, particularly in the anoxic environment of the human gut where oxygen-dependent L-carnitine-metabolizing enzymes are likely inactive. Here, we elucidate the chemical and genetic basis for anaerobic TMA generation from the L-carnitine-derived metabolite γ-butyrobetaine (γbb) by the human gut bacterium *Emergencia timonensis*. We identify a set of genes upregulated by γbb and demonstrate that the enzymes encoded by the induced γbb utilization (*bbu*) gene cluster convert γbb to TMA. The key TMA-generating step is catalyzed by a previously unknown type of TMA-lyase enzyme that utilizes a flavin cofactor to catalyze a redox neutral transformation. We identify additional cultured and uncultured host-associated bacteria that possess the *bbu* gene cluster, providing insights into the distribution of anaerobic γbb metabolism. Lastly, we present genetic, transcriptional, and metabolomic evidence that confirms the relevance of this metabolic pathway in the human gut microbiota. These analyses indicate that the anaerobic pathway is a more substantial contributor to TMA generation from L-carnitine in the human gut than the previously proposed aerobic pathway. The discovery and characterization of the *bbu* pathway provides the critical missing link in anaerobic metabolism of L-carnitine to TMA, enabling investigation into the connection between this microbial function and human disease.

**SIGNIFICANCE:** Trimethylamine (TMA) is a disease-associated metabolite produced in the human body exclusively by microbes. Gut microbes generate TMA from essential nutrients consumed in the human diet, including L-carnitine. However, our understanding of the biochemical mechanisms involved in these transformations is incomplete. In this work, we define the biochemical pathway and genetic components in gut bacteria required for anaerobic production of TMA from γ-butyrobetaine, a metabolite derived from L-carnitine. This discovery identifies a new type of TMA-producing enzyme and fills a critical gap in our knowledge of L-carnitine metabolism to TMA in the anaerobic environment of the human gut. This knowledge will enable evaluation of the link between L-carnitine metabolism and human disease, and the design of potential therapeutics.

## INTRODUCTION

The human gut microbiota collectively synthesizes an array of small molecule metabolites. The metabolic output of this microbial community varies substantially between individual human subjects, and specific metabolites are strongly associated with health and disease (1-3). In many cases, however, we lack both a molecular understanding of how gut microbial metabolites influence human physiology and how the metabolites themselves are produced. These gaps in knowledge limit our ability to establish causative effects of microbial metabolites in human disease and to develop microbiota-based strategies to improve human health. Identification of the specific organisms, genes, and enzymes responsible for metabolite production is needed to accurately profile specific metabolic functions in microbial communities, to experimentally investigate links to disease, and to modulate the metabolic output of the gut microbiota.

Trimethylamine (TMA) is a gut microbial metabolite that has been strongly associated with human disease. It is derived from gut microbial transformations of dietary nutrients including phosphatidylcholine, choline, L-carnitine, betaine, and trimethylamine *N*-oxide (TMAO) (4-9). Microbially produced TMA is absorbed by the host in the gastrointestinal tract, enters hepatic circulation, and is oxidized to TMAO in the liver by the flavin-dependent monooxygenase FMO3 (10). Genetic mutations in the human *FMO3* gene lead to accumulation of TMA in the body, causing the metabolic disorder trimethylaminuria or fish malodor syndrome (11). In addition, elevated plasma levels of TMA and TMAO have been associated with multiple human diseases, including cardiovascular, chronic kidney, and non-alcoholic fatty liver diseases, obesity and type II diabetes, and colorectal cancer (12). Especially strong correlations between TMAO and its precursors have been demonstrated for cardiovascular disease. Elevated plasma levels of dietary TMA precursors were also associated with disease risk, but only when co-occurring with elevated TMAO levels (6, 7, 9). Furthermore, administration of the TMA precursors phosphatidylcholine, choline, and carnitine to atherosclerosis-prone mice resulted in development of atherosclerotic plaques in a gut microbiota-dependent fashion (6, 7). Direct oral administration of TMAO in these mice similarly resulted in phenotypes of atherosclerosis (6). These observations suggest a causal role for TMAO in animal models of cardiovascular disease and that gut microbial metabolism is a crucial factor contributing to pathogenesis. However, a causative role of TMA or TMAO in the development or exacerbation of complex diseases has not yet been definitively established in humans. Deciphering the contribution of TMA production to human disease clearly necessitates a better understanding of the gut microbial metabolic pathways that generate this small molecule.

L-Carnitine is an important dietary precursor to TMA, and its metabolism by gut microbes is associated with cardiovascular disease (13). An essential nutrient for the host, L-carnitine plays a key role in fatty acid β-oxidation by transporting fatty acids across the mitochondrial membrane for metabolism (5, 14). Although it is produced endogenously, humans must uptake additional L-carnitine in the diet to support cellular function (5, 14). The major sources of L-carnitine are animal-based products, especially red meat (15), but it is also ingested as a supplement for enhanced physical performance (16, 17). Early studies of L-carnitine metabolism in rats and human subjects demonstrated its conversion to TMA, the former showing dependence on the gut microbiota (18, 19). More recent studies in humans confirmed the involvement of the gut microbiota in this transformation (7). These studies also noted accumulation of an intermediate metabolite identified as γ-butyrobetaine (γbb) that was also produced by the gut microbiota (18-20). Furthermore, γbb was shown to be a pro-atherogenic metabolite in mouse models like its precursor L-carnitine (20). The well-characterized metabolic pathway that converts L-carnitine to γbb is encoded by the *cai* gene operon (**Figure 1**) and is used during anaerobic respiration by facultative anaerobic Proteobacteria such as *Escherichia coli, Salmonella typhimurium*, and *Proteus mirabilus* (5). The microbial genes and enzymes that are responsible for generating TMA from L-carnitine-derived γbb, however, are not fully elucidated.

**Figure 1.**
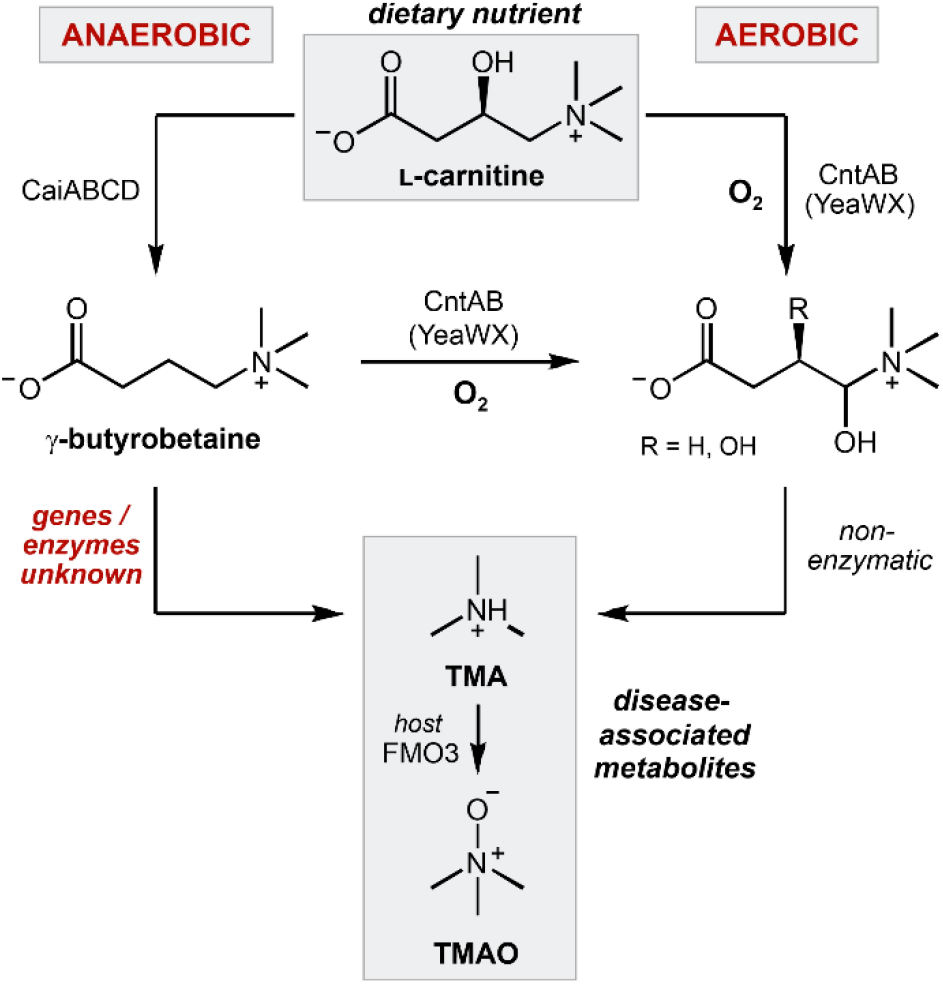
Anaerobic and aerobic bacterial metabolic pathways producing TMA from L-carnitine.

Specifically, there is a significant gap in our understanding of the molecular basis for TMA production from L-carnitine and γbb precursors under anaerobic conditions. Select facultative anaerobic Proteobacteria and Actinobacteria possess an iron-dependent Rieske-type monooxygenase (CntA) that can directly convert carnitine to TMA (20-22). This enzyme uses dioxygen to hydroxylate L-carnitine at the C4 position, followed by non-enzymatic formation of an aldehyde through elimination of TMA (**Figure 1**). Whereas L-carnitine was the only substrate tested for activity with CntA from *Acinetobacter baumannii* (21), the *E. coli* homolog, also known as YeaW (71% amino acid identity), was shown to produce low levels of TMA from both L-carnitine and γbb (20). Although CntA activity was originally proposed to represent the major mechanism for conversion of L-carnitine to TMA by the human gut microbiota, this conclusion has been called into question. This enzyme strictly requires dioxygen for catalysis, however, dioxygen levels in the colon lumen are <1 mm Hg (23). In addition, a study in humans showed that plasma TMAO levels after a carnitine challenge were not correlated with *cntA* gene abundance in gut microbiomes (24). Notably, TMA production from L-carnitine and γbb was demonstrated in anaerobic *ex vivo* incubations of mice cecal and colon tissues (20). Finally, the human gut isolate *Emergencia timonensis*, an obligate anaerobe that does not encode a CntA homolog, was recently found to metabolize γbb to TMA under strictly anaerobic growth conditions (13). Collectively, this information indicates the existence of an as-yet-uncharacterized anaerobic pathway for γbb metabolism in the human gut microbiota.

We present here the identification and characterization of the metabolic pathway, genes, and enzymes responsible for anaerobic TMA production from γbb in *E. timonensis*. The enzyme catalyzing the key C–N bond cleavage reaction that generates TMA is a flavin-dependent, acyl-CoA dehydrogenase-like enzyme that uses the activated CoA thioester of γbb as its substrate. This chemically challenging reaction generates the intermediate crotonyl-CoA, which is further metabolized by *E. timonensis* for anaerobic respiration and as a source of carbon and energy. Homologous gene clusters for γbb metabolism are present in other cultured and uncultured host-associated bacteria from the Clostridiales order. We find that anaerobic γbb metabolism is prevalent in human gut microbiomes and is likely a major, underappreciated contributor of L-carnitine-derived TMA. Together, this work expands our knowledge of TMA-producing enzymes, pathways, and organisms, providing a more complete understanding of microbial TMA production in the anoxic human gut. These findings identify potential new targets for manipulation of this microbial function and will help resolve the major dietary and microbial contributors to TMA production.

## RESULTS

### γ-Butyrobetaine induces expression of a candidate gene cluster in *E. timonensis* SN18

Only a single cultured bacterium, a human fecal isolate of *E. timonensis*, has been reported to produce TMA from γbb under anaerobic conditions (13). We began our efforts to identify the genes and enzymes responsible for γbb metabolism by testing the type strain *E. timonensis* SN18 for this activity. Indeed, anaerobic cultures of *E. timonensis* SN18 supplemented with γbb demonstrated complete consumption of this substrate and production of a stoichiometric amount of TMA (**Figure S1**). When the cultures were supplemented with [*N*-(CD_3_)_3_]-γbb, deuterium-labeled TMA was detected by liquid chromatography–tandem mass spectrometry (LC–MS/MS) (**Figure S1**), confirming its origin from γbb. Resting cell suspensions of *E. timonensis* SN18 also fully converted γbb to TMA (**Figure 2A**). However, this activity was only observed when the cells had been cultured in medium containing γbb; resting cell suspensions of *E. timonensis* SN18 cultured in the absence of γbb were unable to consume γbb and did not generate TMA (**Figure 2A**). Together, these results suggest that γbb induces expression of the genes involved in its metabolism.

**Figure 2.**
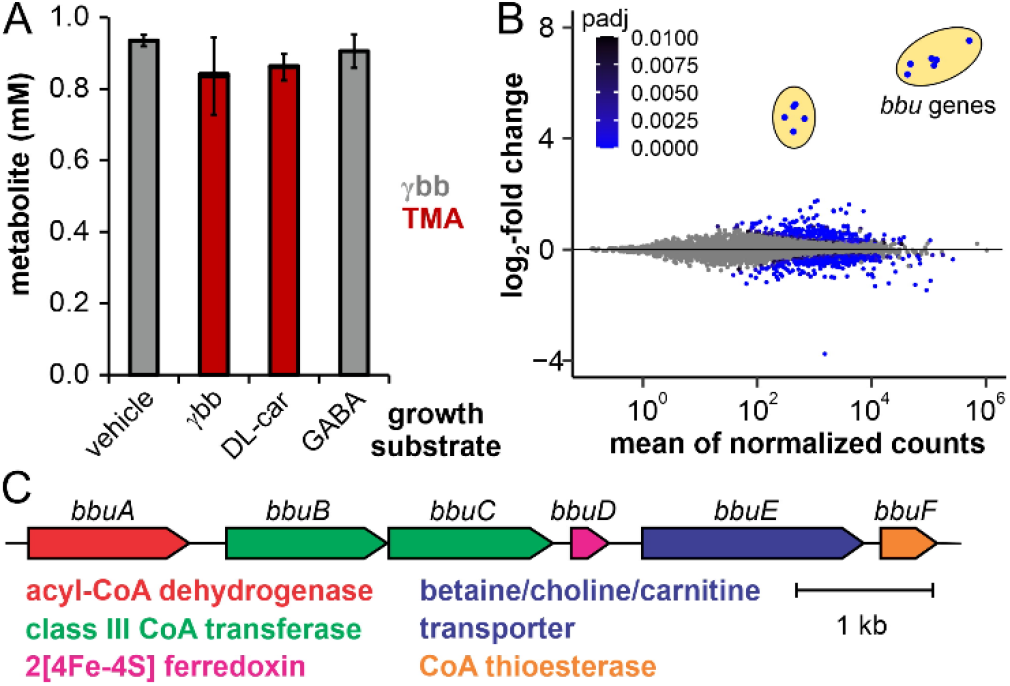
γbb induces expression of the *bbu* gene cluster in *E. timonensis* SN18. (A) Stacked bar plots of γbb and TMA concentrations after a 3 h incubation of 1 mM γbb in resting cell suspensions of *E. timonensis* SN18 previously cultured in media supplemented with 1 mM γbb, DL-carnitine, GABA, or 1× PBS (vehicle). Error bars represent the standard deviation from the mean of three biological replicates. (B) Differential gene expression from *E. timonesis* SN18 cultures supplemented with γbb or treated with a vehicle at an OD = 0.7, plotted against the mean of normalized counts from three biological replicates. The two sets of differentially upregulated genes are circled in yellow. Genes with an adjusted *p*-value >0.01 (Wald test) comparing γbb- and vehicle-induced cells are represented by grey circles. (C) The *bbu* gene cluster and automated protein function annotations.

We also evaluated the ability of *E. timonensis* SN18 to metabolize DL-carnitine and γ-aminobutyric acid (GABA), close structural homologs of γbb. Neither of these substrates was consumed during growth (**Figure S1**) nor in resting cell suspensions of cultures supplemented with γbb (**Figure S1**). Next, we tested whether these substrate analogs could induce expression of the γbb metabolic enzymes. Resting cell suspensions of cultures supplemented with GABA did not exhibit γbb metabolism; however, resting cell suspensions from cultures supplemented with DL-carnitine did convert γbb to TMA (**Figure 2A**). Thus, while DL-carnitine appears to induce expression of the genes responsible for γbb metabolism in *E. timonensis*, the metabolic enzymes are unable to use DL-carnitine as a substrate for TMA generation.

The inducible metabolism of γbb in *E. timonensis* SN18 suggested that the genes involved in this pathway could be discovered using RNA-sequencing (RNA-seq). Only two sets of genes were both highly and differentially expressed in γbb-induced cultures compared to control cultures grown in the absence of γbb (**Figure 2B, Dataset S1**). As expected from our previous cell-based activity experiments, expression of these genes was also upregulated in cells grown with DL-carnitine, but not with GABA (**Figure S2, Dataset S1**). The most differentially upregulated genes are colocalized in the *E. timonensis* SN18 genome within a region we designate the γ-<u>b</u>utyro<u>b</u>etaine <u>u</u>tilization (*bbu*) gene cluster (**Figure 2C**). This six-gene cluster encodes a predicted flavin-dependent acyl-CoA dehydrogenase (*bbuA*), two class III CoA-transferases (*bbuB* and *bbuC*), a di-[4Fe-4S]-cluster ferredoxin (*bbuD*), a betaine/carnitine/choline transporter (*bbuE*), and an acyl-CoA thioesterase (*bbuF*). The second set of genes upregulated in γbb-induced cells encodes for predicted riboflavin biosynthesis enzymes (**Dataset S1**), which likely support cofactor biosynthesis for the enzyme encoded by *bbuA*. The highly specific transcriptional response to γbb suggested that the *bbu* gene cluster was involved in γbb metabolism. In addition, the presence of a gene encoding a transporter for trimethylammonium-containing compounds (*bbuE*) was consistent with this hypothesis.

### The CoA transferases BbuB/C initiate γbb metabolism via formation of a γbb-CoA thioester

Based on the predicted annotations of the *bbu* genes, we hypothesized that γbb would be initially activated for subsequent TMA elimination through formation of a CoA thioester. The conversion of a carboxylic acid to a thioester is a common first step in many metabolic pathways as it lowers the p*K*_a_ of the C_α_-protons, facilitating further reactions (25, 26). Indeed, in resting cell suspensions incubated with γbb, a metabolite was detected by LC–MS with a peak retention time and mass-to-charge ratio (895.5 *m/z*, [M+H]^+^) matching those of a γbb-CoA standard (**Figure 3A, Figure S3**). The origin of this metabolite was confirmed by incubating cell suspensions with deuterium-labeled 2,2,3,3,4,4-D_6_-γbb (D_6_-γbb), which led to an increase of 6 Da (901.5 *m/z*, [M+H]^+^) for the assigned γbb-CoA peak (**Figure 3A, Figure S3**). These results support the hypothesis that γbb metabolism proceeds via a CoA thioester intermediate.

**Figure 3.**
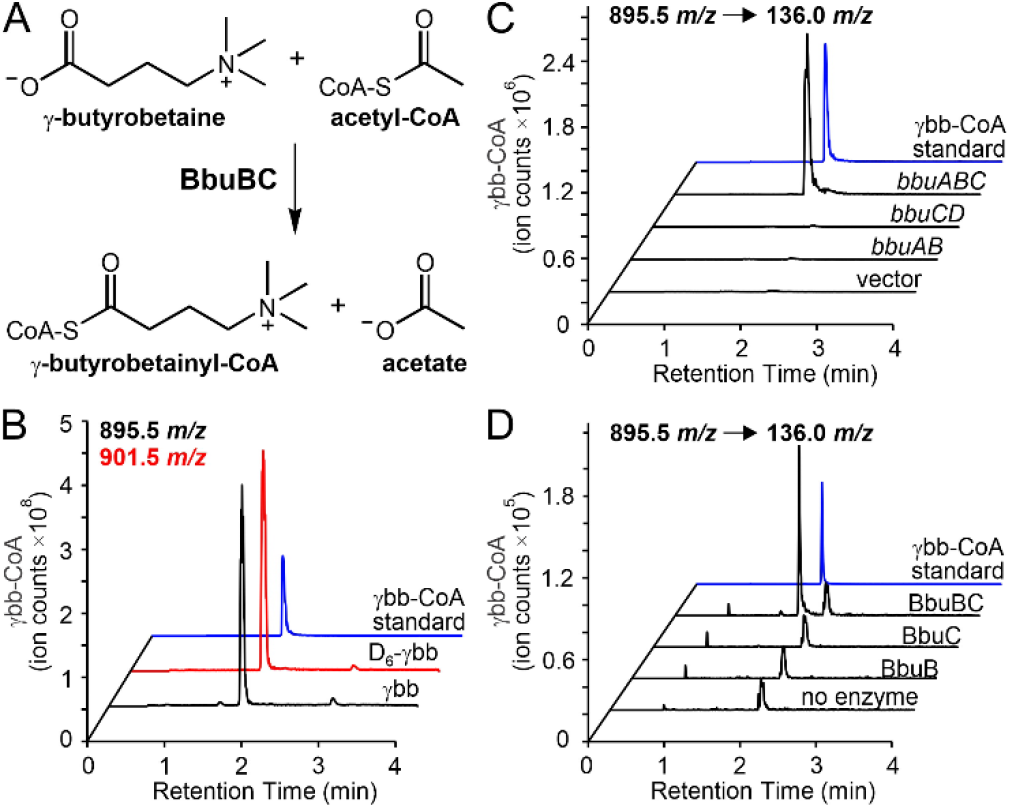
γbb-CoA is the first intermediate in γbb metabolism. (**A**) Chemical reaction catalyzed by the BbuB and BbuC proteins. (**B**) LC–MS extracted ion chromatograms (EIC) of γbb-CoA ([M+H]^+^ = 895.5 *m/z*) and D_6_-γbb-CoA ([M+H]^+^ = 905.5 *m/z*) from extracts of *E. timonensis* cell suspensions incubated for 40 min with γbb (black) or D_6_-γbb (red), respectively, compared to a γbb-CoA standard (blue). (**C**) LC–MS/MS selected ion chromatograms (SIC) of the 136.0 *m/z* fragment ion of γbb-CoA produced from a 1 h incubation of γbb and acetyl-CoA with crude lysate of *E. coli* constitutively expressing *bbu* genes or empty vector. (**D**) LC–MS/MS SIC of the 136.0 *m/z* fragment ion of γbb-CoA produced from a 1 h incubation of γbb and acetyl-CoA with purified recombinant proteins. **Figure S3** shows relative amounts of γbb-CoA from three biological replicates of experiments described for panels C and D.

We next investigated whether two CoA transferases encoded by the *bbu* gene cluster, BbuB and BbuC, were directly involved in γbb-CoA production. Indeed, γbb-CoA was detected from incubations of γbb with crude lysate of *E. coli* cells constitutively expressing the first three genes of the cluster (*bbuABC*) (**Figure 3B**). Interestingly, γbb-CoA was not detected when either of the two CoA transferase genes was expressed alone (**Figure 3B**). Production of γbb-CoA was also observed when recombinant, purified BbuB and BbuC were added together with γbb to lysate of *E. coli* transformed with empty vector (**Figure S3**), but only when both proteins were present. The requirement for both CoA transferase proteins was confirmed *in vitro*, suggesting that the two proteins may form an active complex. In these experiments, the BbuB and BbuC proteins together generated γbb-CoA from γbb and an appropriate acyl-CoA donor substrate (**Figure 3C**). Various short-chain fatty acyl-CoAs (acetyl-, butyryl-, propionyl-, and crotonyl-CoA) were tested as CoA-donating substrates in the reaction, and while all compounds resulted in production of γbb-CoA, acetyl-CoA was the preferred co-substrate (**Figure S3**). We also tested DL-carnitine and GABA as alternative CoA acceptors *in vitro* with BbuB/C and acetyl-CoA as the donor. However, neither of the presumed products, carnitinyl-CoA or γ-aminobutyryl-CoA, was detected by LC–MS/MS after a 3 h incubation (**Figure S3**). The high selectivity of the CoA transferases BbuB and BbuC for γbb directly connects the *bbu* gene cluster to γbb metabolism.

### The flavoenzyme BbuA catalyzes elimination of TMA from γbb-CoA

The key step in TMA production from γbb is breaking an unactivated C–N bond. None of the *bbu* genes is predicted to encode homologs of known enzymes that catalyze C–N bond cleavage and release TMA (*e*.*g*. GRE choline TMA-lyase (27), glycine betaine reductase (28), ergothionase (29)), indicating that this pathway uses a distinct enzyme for this critical step. We identified the predicted acyl-CoA dehydrogenase-like enzyme BbuA as the most likely candidate for this reaction. Enzymes belonging to the acyl-CoA dehydrogenase family are flavin-dependent oxidoreductases that act on acyl-CoA thioester substrates, typically installing or removing an α,β-unsaturation through two-electron oxidation or reduction reactions, respectively (30). We envisioned that BbuA might instead use its flavin cofactor in a cryptic radical mechanism to activate γbb-CoA for TMA elimination from the C4 position. This proposed reaction resembles the C4 elimination of water from 4-hydroxybutyryl-CoA catalyzed by the FAD-dependent enzyme 4-hydroxybutyryl-CoA dehydratase (4HBD) involved in bacterial GABA and succinate metabolism (31). While BbuA and 4HBD are predicted to have a similar structure and flavin cofactor, they share very minimal sequence identity (∼10%).

Initial attempts to access soluble BbuA protein via heterologous expression in *E. coli* were unsuccessful. Soluble BbuA protein was only obtained under the following conditions: 1) co-expression with the native GroEL-ES proteins from *E. timonensis* SN18, 2) addition of exogenous riboflavin to the growth medium, and 3) addition of excess FAD during cell lysis. Crude lysates of *E. coli* cells over-expressing the *bbuABC* genes under these conditions demonstrated TMA production when incubated anaerobically with γbb and acetyl-CoA (**Figure S4**). Purified BbuA was then tested for activity *in vitro* when incubated with γbb, acetyl-CoA, and the CoA transferases BbuBC. Robust TMA production was observed after a 1 h incubation without redox mediators (**Figure 4B**), demonstrating that these reaction components are sufficient to catalyze multiple turnovers of γbb. Finally, γbb-CoA was tested directly as the substrate for BbuA activity *in vitro*. This reaction also resulted in TMA production (**Figure 4C**). These results conclusively demonstrate that BbuA is the critical TMA-generating enzyme in anaerobic γbb metabolism.

**Figure 4.**
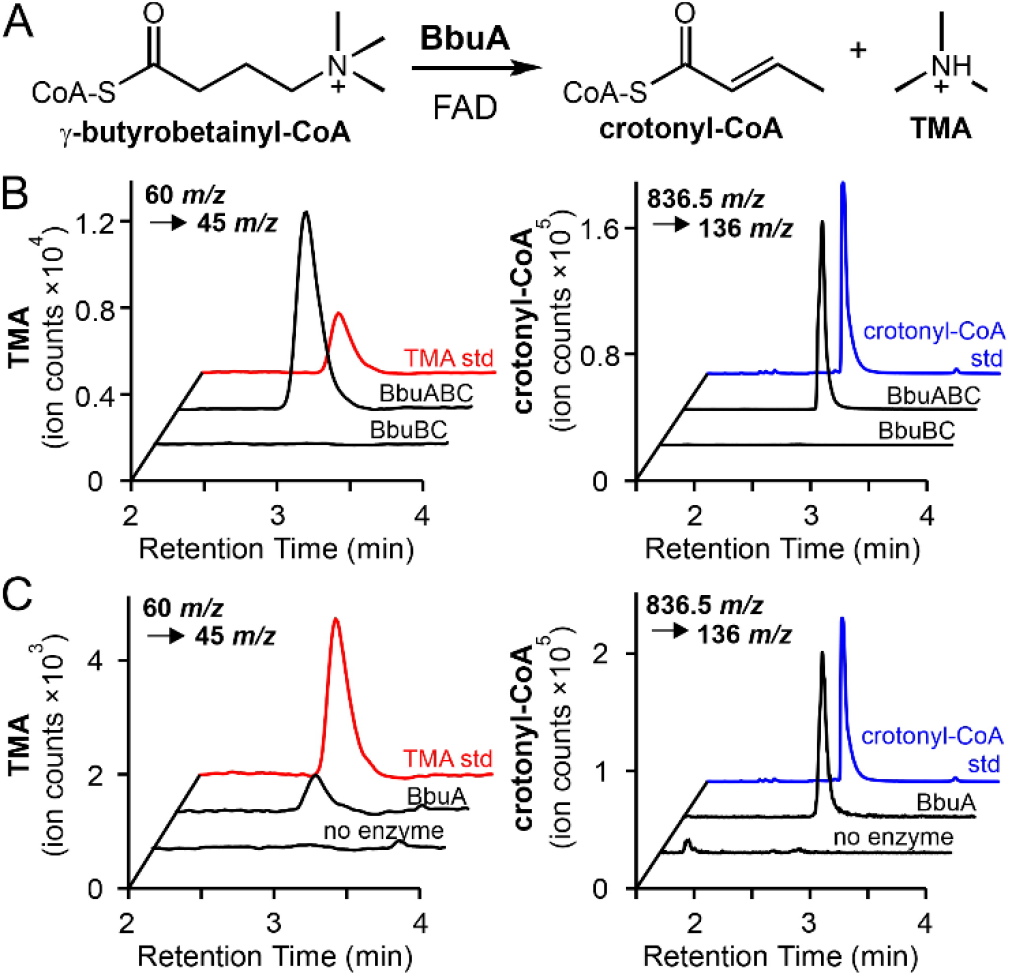
The FAD-dependent enzyme BbuA generates TMA. (**A**) Chemical reaction catalyzed by BbuA. (**B**) LC–MS/MS SIC of the precursor-fragment ion pairs of TMA (left) and crotonyl-CoA (right) produced from 1 h reactions containing γbb, acetyl-CoA, BbuB, BbuC, and FAD with or without addition of BbuA. (**C**) LC–MS/MS SIC of the precursor-fragment ion pairs of TMA (left) and crotonyl-CoA (right) produced from 1 h reactions containing γbb-CoA and FAD with or without addition of BbuA. **Figure S4** shows quantification of TMA concentration and relative amounts of crotonyl-CoA from three biological replicates of experiments described for panels B and C.

Next, we sought to determine the co-product of the BbuA-catalyzed reaction. We predicted that crotonyl-CoA would be generated upon elimination of TMA from γbb-CoA by analogy to 4-hydroxybutyryl-CoA dehydration in GABA/succinate metabolism (31). Indeed, crotonyl-CoA was detected by LC–MS/MS in both the *E. coli* crude lysate incubations (**Figure S4**) and the *in vitro* assays with purified BbuA (**Figure 4**). Both the retention time and *m/z* of the detected metabolite were consistent with a crotonyl-CoA standard (**Figure S4**). The production of crotonyl-CoA supports the proposed redox-neutral TMA elimination from γbb-CoA (**Figure 4**). Together, the gain-of-function experiments in *E. coli* and *in vitro* activity assays demonstrate that the *bbu* gene cluster is responsible for TMA production from γbb, and that the three genes *bbuABC* are necessary and sufficient to confer this function in *E. coli* and *in vitro*.

### *E. timonensis* metabolizes γbb for anaerobic respiration and carbon acquisition

Although crotonyl-CoA was produced in lysate and *in vitro* reactions containing BbuA, crotonyl-CoA did not accumulate in resting cell suspensions of *E. timonensis* SN18 incubated with γbb. Instead, we detected the known products of bacterial crotonyl-CoA metabolism: 3-hydroxybutyryl-CoA, acetyl-CoA, and butyryl-CoA (**Figure 5B**). Inspection of their mass spectra showed deuterium incorporation when D_6_-γbb was used, confirming their origin from γbb (**Figure S5**). These results suggest that crotonyl-CoA generated from γbb is further metabolized by *E. timonensis*. Notably, the *bbu* gene cluster does not encode any enzymes known to be involved in crotonyl-CoA metabolism (32). However, protein homologs are encoded by constitutively expressed genes located elsewhere in the *E. timonensis* SN18 genome (**Dataset S2**). To test whether they could account for the observed crotonyl-CoA metabolism, crotonyl-CoA was incubated with cell lysates of *E. timonensis* SN18 grown in the presence or absence of γbb. In both cases, crotonyl-CoA was converted to 3-hydroxybutyryl-CoA during an hour incubation (**Figure S6**). Longer incubations led to accumulation of acetyl-CoA and low levels of butyryl-CoA (**Figure S6**). These results demonstrate that the final steps of γbb metabolism in *E. timonensis* SN18 are known transformations of crotonyl-CoA, catalyzed by constitutively produced enzymes encoded outside of the *bbu* gene cluster (**Figure 5A**).

**Figure 5.**
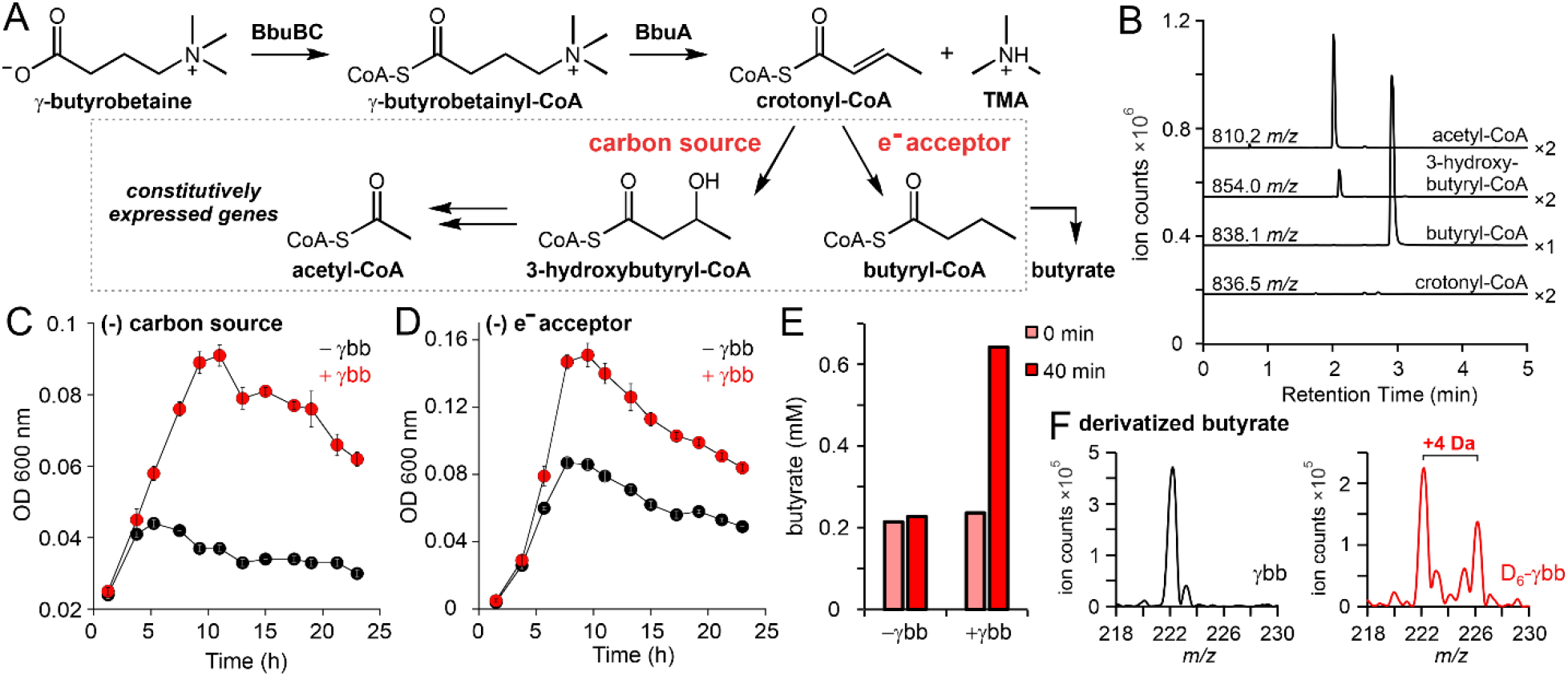
γbb-derived crotonyl-CoA is further metabolized by *E. timonensis* for carbon and respiration. (**A**) Complete metabolic pathway using γbb as a carbon source and electron acceptor.(**B**) LC–MS/MS SICs of parent ions with 136.0 *m/z* fragments that correspond to expected products of crotonyl-CoA metabolism from extracts of *E. timonensis* SN18 cell suspensions incubated with γbb for 40 min. Chromatogram magnification is indicated next to each trace. (**C-D**) Anaerobic growth curves of *E. timonensis* SN18 in minimal media lacking (**E**) carbon sources or (**F**) electron acceptors with (red) or without (black) γbb. Error bars represent standard deviation from the mean of three biological replicates. (**E**) Concentrations of derivatized butyrate detected from extracts of *E. timonensis* cell suspensions incubated for 0 (pink) or 40 (red) min in the presence or absence of 1 mM γbb. Bars show the average of two biological replicates. (**F**) Mass spectra of derivatized extracts (unlabeled butyrate [M+H]^+^ = 222 *m/z*) of *E. timonensis* cell suspensions incubated for 40 min with γbb (black) or D_6_-γbb (red).

Some species of Clostridia can use crotonyl-CoA metabolism for anaerobic respiration via reduction to butyryl-CoA, and as a carbon and energy source through conversion to acetyl-CoA (**Figure 5A**) (33, 34). We examined whether γbb metabolism via crotonyl-CoA would enhance the growth of *E. timonensis* SN18 in media conditions with limited electron acceptors or carbon sources. In both culture conditions, the growth yield was markedly enhanced with γbb supplementation (**Figure 5C, 5D**) and the growth rate was consistent with the rates of γbb depletion and TMA production (**Figure S7**). These results support a key role for crotonyl-CoA as an intermediate in γbb metabolism and indicate a physiological role for this activity in *E. timonensis*.

Due to its prominent roles in the human gut, we next investigated whether free butyrate was produced from *E. timonensis* γbb metabolism. Using a carboxylate-derivatization method and LC–MS, elevated levels of butyrate were detected in suspensions of γbb-induced cells incubated with γbb compared to suspensions of non-induced cells (**Figure 5E, Figure S8**). In addition, deuterium incorporation into butyrate was observed when D_6_-γbb was used (**Figure 5F**). Analysis of the mass spectrum revealed that the D_6_-isotopolog was not present; rather the +4 Da species was the primary isotopolog detected (**Figure 5F, Figure S8**). The loss of two deuteria in the butyrate product is consistent with a crotonyl-CoA precursor. To rule out the possibility that the loss of label resulted from exchange of the acidic C_α_-deuterons with protons from solution, a control experiment was performed incubating perdeuterated D_7_-butyrate with cell suspensions. The mass spectrum of butyrate from this experiment showed minimal deuterium exchange with solvent protons during the same incubation period (**Figure S8**). In addition, γbb-CoA detected from incubations with D_6_-γbb showed minimal deuterium loss (**Figure S8**). These control reactions support the proposal that γbb metabolism in *E. timonensis* proceeds through the α,β-unsaturated crotonyl-CoA intermediate and generates free butyrate, an important metabolite in the human gut environment.

### The *bbu* gene cluster predicts γbb metabolism in bacterial isolates

As noted previously, *E. timonesis* is the only bacterial species known to metabolize γbb to TMA under anaerobic conditions. Having discovered the gene cluster responsible for this activity, we searched for homologous gene clusters in other sequenced bacterial genomes. We identified *bbu*-like gene clusters in four host-associated, cultured bacteria belonging to the order Clostridiales. To examine whether the *bbu* genes are diagnostic of γbb metabolism, two commercially available strains, the human oral isolate *Eubacterium minutum* ATCC 700079 and the feline gut isolate *Agathobaculum desmolans* ATCC 43058 (previously *Eubacterium desmolans* or *Butyricoccus desmolans*), were tested for this activity. Both strains converted γbb to TMA in anaerobic cultures (**Figure 6A**), demonstrating that the presence of the *bbu* gene cluster predicts the capacity for anaerobic γbb metabolism. The rarity of this gene cluster in sequenced genomes of cultured bacteria motivated us to search uncultured bacterial genomes that had been assembled from human metagenomes (*i*.*e*. metagenome-assembled genomes, MAGs) (35, 36). We identified homologous *bbu* gene clusters in 11 uncultured, human-associated MAGs, all of which belong to the order Clostridiales (**Figure 6B, Dataset S3**). Two of the uncultured bacterial genomes were assembled from human oral samples, while the remaining genomes harboring the *bbu* gene cluster were assembled from human stool samples. The presence of the *bbu* gene cluster in multiple uncultured gut bacteria suggests that γbb metabolism is underrepresented among cultured isolates and that *E. timonensis* may not be solely responsible for this metabolism in the human gut. Finally, we searched for the presence of the *cai* genes responsible for metabolizing L-carnitine to γbb in these *bbu*-containing bacterial genomes. However, none of these genomes possess homologs of known *cai* genes, indicating that these γbb-metabolizing bacteria are incapable of the full conversion of L-carnitine to TMA.

**Figure 6.**
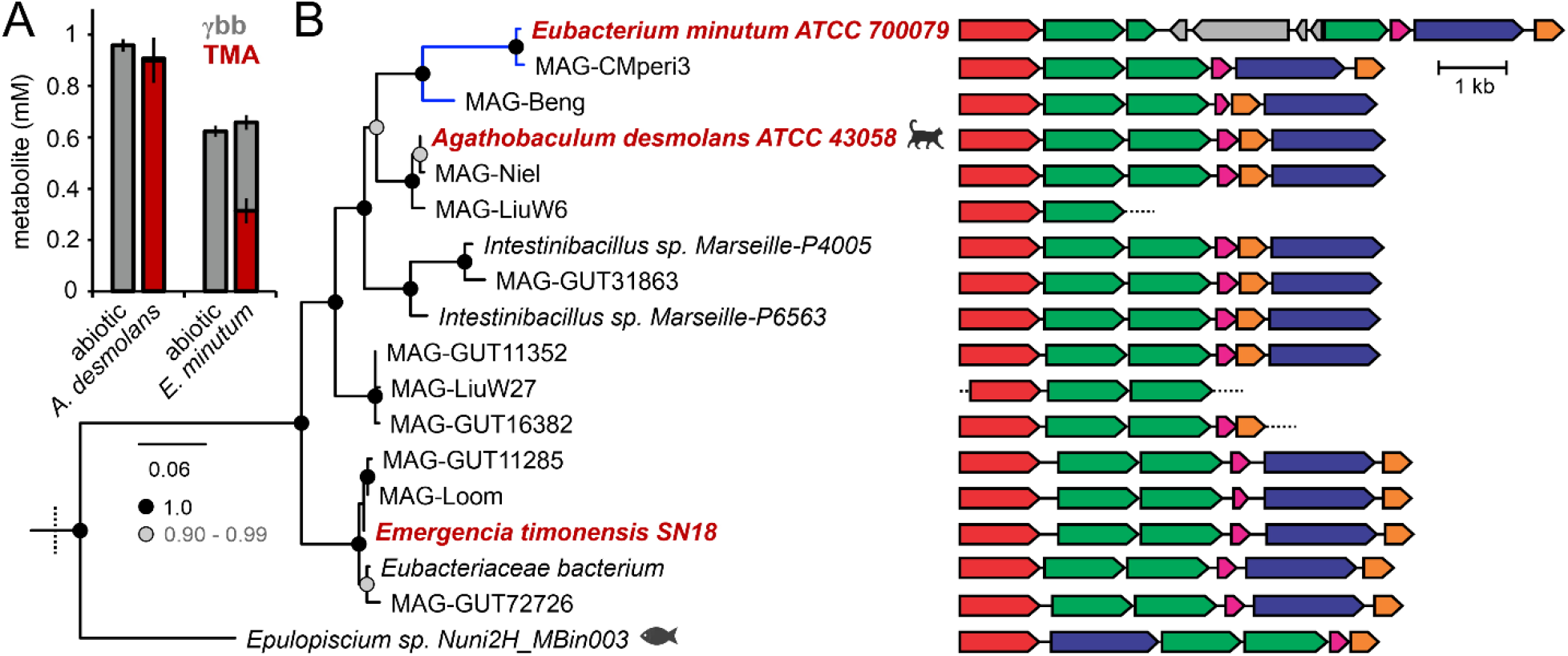
Cultured and uncultured host-associated Clostridiales bacteria possess *bbu*-like gene clusters indicative of γbb metabolism. (**A**) TMA and γbb detected by LC–MS/MS from pure cultures of *A. desmolans* and *E. minutum* grown in rich media supplemented with γbb. Error bars represent the standard deviation from the mean of three biological replicates. (**B**) BbuA clade of a maximum likelihood phylogenetic tree (**Figure Sx**) and genomic context of BbuA homologs. aBayes branch supports are shaded based on the legend. Red labels indicate experimentally validated γbb-metabolizing bacteria. Blue branches indicate an isolate or MAG originating from an oral sample.

### Presence of *bbu* genes in human gut metagenomes is correlated with lower γbb levels

Motivated by the presence of the *bbu* gene cluster in uncultured gut bacteria and the implications for γbb metabolism in human health, we assessed the prevalence of the *bbu* gene cluster in publicly available human metagenomic datasets. Our analysis used the *bbuA* gene as a representative of the full cluster in stool metagenomes from healthy adult populations. We detected hits for the *bbuA* gene in samples from every cohort, but the proportion of *bbuA*-positive samples varied substantially between cohorts, ranging from 15–85% positive (**Figure 7A**). We also quantified the abundance of the *bbuA* gene in *bbuA*-positive samples and found that it was consistent across cohorts at 1:1000 genes per microbial genome, with a range of 1:10^2^ to 1:10^4^ genes per microbial genome (**Figure S9**). Since diet has been reported as an important factor in TMA production capacity of the microbiota, we examined whether *bbuA* gene presence or abundance was correlated with diet. However, neither *bbuA* gene presence nor abundance was correlated with self-reported omnivorous or vegetarian donors from the BIO-ML cohort (**Figure S10**). Overall, the distribution profile of *bbu* genes suggests that γbb-metabolizing bacteria are frequently present in the human gut microbiota but are low-abundance members of the community.

**Figure 7.**
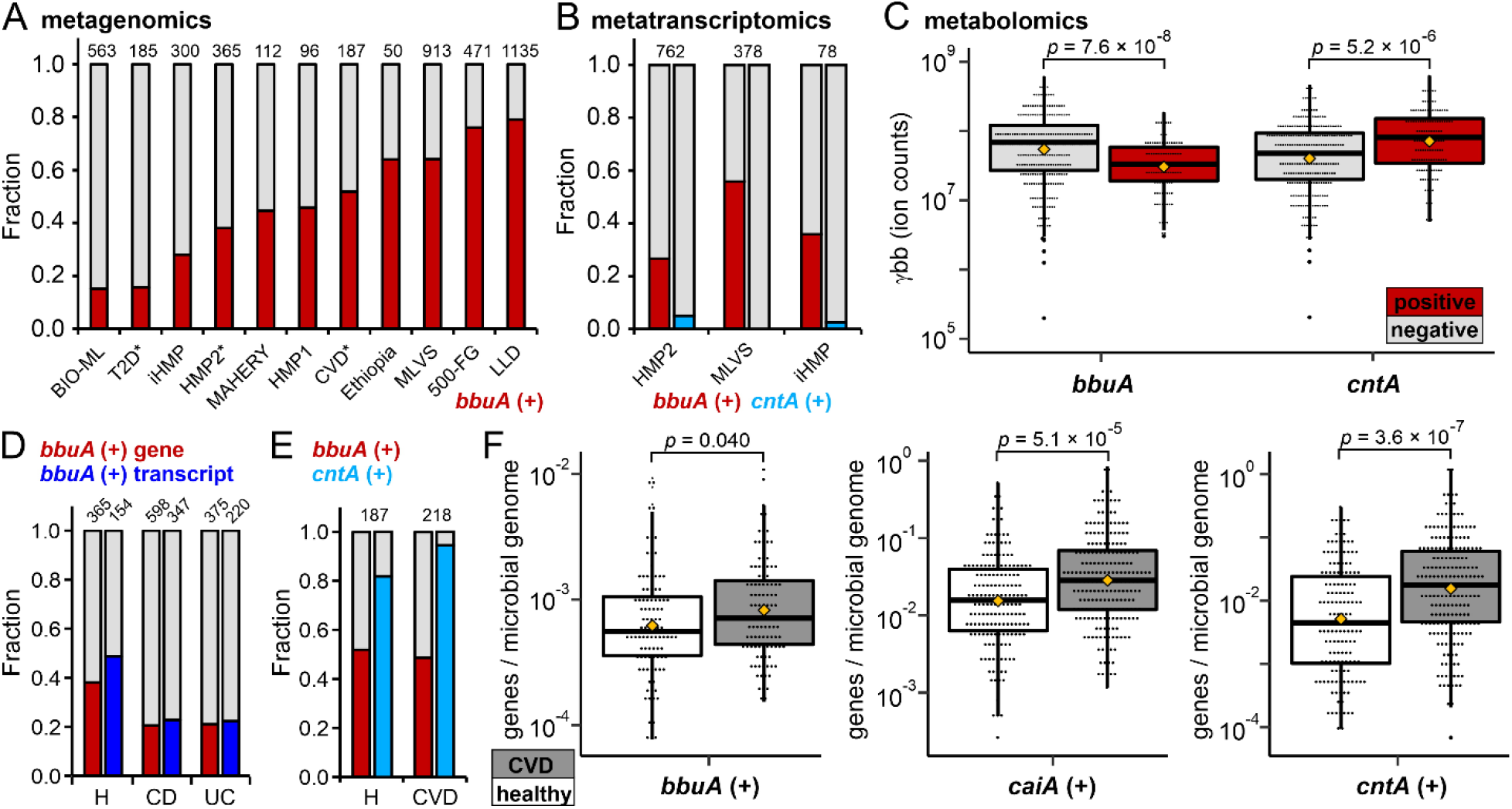
Anaerobic γbb metabolism genes are widely distributed and expressed in the human gut microbiota. (**A**) Fraction of stool metagenomes that are positive (red) and negative (grey) for the *bbuA* gene from healthy donors in human studies (**Dataset S4**). Study sample size (*N*) is indicated above each bar. Asterisks indicate that only healthy donors from those projects were included in this analysis. (**B**) Fraction of stool metatranscriptomes that are positive and negative for *bbuA* or *cntA*. Study sample size (*N*) is indicated above each bar. (**C**) Correlation between the presence of *bbuA* or *cntA* genes in metagenomes and γbb metabolite levels in HMP2 stool samples. Mean values are represented by yellow diamonds and *p*-values were determined using the Mann-Whitney *U*-test. (**D**) Fraction of samples positive for the *bbuA* gene or transcripts in HMP2 IBD cohorts compared to healthy controls. Study sample size (*N*) is indicated above each bar. (**E**) Fraction of samples positive for the *bbuA* or *cntA* gene in a CVD cohort compared to healthy controls. Study sample size (*N*) is indicated above each bar. (**F**) Correlations between the presence of the *bbuA, caiA*, or *cntA* genes and CVD compared to healthy controls. Mean values are represented by yellow diamonds and *p*-values were determined using the Mann-Whitney *U*-test.

To provide further support for the relevance of the *bbu* pathway in the human gut, we evaluated gene expression using metatranscriptomic data from three cohorts – HMP2 (37), iHMP (38), and MLVS (39). We detected *bbuA* transcripts in 27–56% of samples in these cohorts (**Figure 7B**), suggesting that the *bbu* pathway is expressed in the human gut. Transcripts of the *bbuA* gene were detected in samples that are positive for the *bbuA* gene, but also a subset (16–26%) of *bbuA*-negative samples (**Figure S11**). This observation suggests that the *bbuA* gene abundance was too low to be detected by the metagenomic sequencing, but that the transcript levels were high enough to be captured by the read depth of the metatranscriptomic sequencing. Notably, very few samples in these cohorts (0–4%) were positive for *cntA* transcripts (**Figure 7B, Figure S11**), consistent with previously reported analyses (40). Thus, the metatranscriptomic data suggest that the *bbu* pathway is more likely to be functionally operant in the human gut than the O_2_-dependent pathway mediated by *cntA*.

Next, we examined three cohorts with corresponding stool metabolomics data (HMP2 (37), BIO-ML (41), and PRISM (42)) to evaluate whether the presence of *bbuA* gene or transcripts is associated with changes in metabolite levels. In each of the three independent cohorts, the levels of γbb were lower in samples that were positive for the *bbuA* gene compared to those that were negative (**Figure 7C, Figure S12**). In the HMP2 cohort, the samples positive for *bbuA* transcripts were also correlated with lower γbb levels (**Figure S13**). The *bbuA* gene, however, was not associated with stool carnitine or butyrate levels (**Figure S14**). In contrast to *bbuA*, the presence of the *cntA* gene was correlated with higher or equivalent γbb levels compared to *cntA*-negative samples rather than lower (**Figure 6C, Figure S12**). These results suggest that the *bbu* gene cluster may be primarily responsible for γbb consumption in the human gut.

Finally, we evaluated whether differences in the presence or abundance of the *bbuA* gene are correlated with human disease. Although TMA production is not known to be a factor in inflammatory bowel disease (IBD), we noticed that the *bbuA* gene was detected in fewer samples from donors with Crohn’s disease and ulcerative colitis compared to healthy controls in both HMP2 (37) and PRISM (42) cohorts (**Figure 7D, Figure S15**). The same trend was observed when analyzing *bbuA* transcripts in the HMP2 study (**Figure 7D**). However, there was no difference in gene abundance between the IBD and healthy cohorts (**Figure S15**). Next, we analyzed metagenomes of a cardiovascular disease (CVD) cohort from China (43). The proportion of *bbuA*-positive samples was not different between healthy and CVD groups (**Figure 7E**), but the gene abundance in *bbuA*-positive samples was modestly higher in CVD donors compared to healthy controls (**Figure 7F**). A more significant elevation was found in the *caiA* gene abundance in the CVD cohort compared to the healthy group (**Figure 7F**), suggesting that the conversion of L-carnitine to γbb may be a more relevant biomarker for CVD than the *bbu* gene cluster. Consistent with previous reports (40), we also detected a higher prevalence and abundance of the *cntA* gene in CVD samples compared with healthy controls from this study (**Figure 7E-F**). However, the lack of correlation between *cntA* gene presence and expression observed in other cohorts raises questions about the functional relevance of this finding.

## DISCUSSION

Gut microbial production of TMA from dietary L-carnitine has been recognized for decades, but our knowledge of the reactions, enzymes, and organisms involved in the O_2_-independent pathway has remained incomplete. We used RNA-seq to identify a substrate-inducible *bbu* gene cluster that is responsible for anaerobic γbb metabolism in the gut microbe *E. timonensis*. We demonstrated that the enzymes encoded within the *bbu* gene cluster are necessary and sufficient to convert γbb to TMA. With the discovery of the genetic basis for anaerobic γbb metabolism, we then determined the chemical steps involved, identified other gut microbes with this function, and evaluated the relevance of this microbial activity in human populations.

The pathway for anaerobic metabolism of γbb provides an interesting parallel to GABA metabolism by gut microbes (44). Given the structural similarity of the two molecules, which differ by only the *N*-methylation of γbb, the use of comparable pathways for their catabolism is perhaps not surprising. Analogous chemical steps, including substrate activation via CoA formation and C4 elimination, occur in both pathways to yield the common product crotonyl-CoA. This intermediate is then processed by the bacteria as a carbon and energy source or as an electron acceptor. Yet there are notable distinctions in the substrate activation steps and key lyase enzymes of each pathway. Conversion of the carboxylate to the thioester lowers the p*K*_a_ of the C_α_-protons (25, 26), facilitating elimination of the C4 hydroxyl or trimethylammonium group in the 4HBD- or BbuA-catalyzed reaction, respectively. Due to its positive charge, the trimethylammonium moiety of γbb-CoA is a good leaving group for C4 elimination. At physiological pH, the primary amine of GABA would also be protonated and positively charged (45), which could enable an analogous direct elimination from a hypothetical γ-aminobutyryl-CoA intermediate. Interestingly, GABA-metabolizing bacteria instead convert the amine into a hydroxyl group, which is a worse leaving group, prior to formation of the CoA thioester and C4 elimination. This biochemical pathway may be the result of repurposing the 4HBD enzyme from succinate metabolism in bacteria and archaea (46-48), which proceeds via a 4-hydroxybutyryl-CoA intermediate. The flavin-dependent lyase enzymes, 4HBD and BbuA, that catalyze the key C4 elimination step in these pathways both belong to the broad structural superfamily of acyl-CoA dehydrogenases (30). This generic functional annotation implies two-electron redox chemistry using a flavin cofactor. We hypothesize instead that BbuA may use a radical-based mechanism similar to that proposed for 4HBD (49), wherein the flavin cofactor enables single-electron chemistry to mediate TMA elimination. Interestingly, however, these two enzymes do not appear to be closely related homologs based on the negligible sequence identity to one another. Therefore, if 4HBD and BbuA use similar mechanistic strategies, these enzymes may be an interesting example of convergent evolution to achieve a chemically challenging elimination reaction. The mechanism of the TMA-lyase reaction catalyzed by BbuA will be the subject of future studies.

The BbuA enzymatic reaction is unique compared to other known TMA-producing enzymes. The L-carnitine monooxygenase CntA uses a metallocofactor to activate dioxygen for substrate hydroxylation, generating a chemically unstable product that breaks down to eliminate TMA. Conversely, BbuA directly catalyzes the C–N bond cleavage step. The enzyme glycine betaine reductase catalyzes an overall two-electron reduction of glycine betaine to produce TMA, employing protein-substrate covalent adducts via (seleno)cysteine residues (28). In contrast, BbuA catalyzes a redox-neutral transformation and uses an organic flavin cofactor. The choline TMA-lyase CutC is a glycyl radical enzyme which uses protein-based radicals to initiate catalysis (27). CutC is proposed to generate a carbon-centered substrate radical intermediate that undergoes a spin center shift and promotes direct 1,2 elimination of TMA (50). The BbuA enzyme could share a similar radical-based mechanistic logic with CutC but, instead of protein-based radicals, uses a flavin cofactor for radical generation. However, the chemical differences between the γbb and choline substrates – the unactivated carbon chain attached to the trimethylammonium in γbb compared to the vicinal hydroxyl group in choline that actively participates in TMA elimination – will necessitate mechanistic divergence. Intriguingly, the mechanism employed by CutC could theoretically operate using L-carnitine as the substrate, as it also has a vicinal alcohol, yet such a hypothetical reaction is not known to exist.

The metabolic pathways that produce TMA from both choline and γbb are used by gut bacteria to access a source of carbon and energy; in addition, γbb metabolism is a mode of anaerobic respiration. This latter pathway is yet another example of reductive chemistry performed by microbes in the anaerobic gut environment. The product of γbb respiration is the short-chain fatty acid butyrate, which has many established roles in host-microbiota interactions (51). It is a major source of energy for intestinal epithelial cells (52) and is an important signaling molecule as a GPCR ligand (51). The physiological roles of butyrate generally promote host health, for example, by enhancing gut barrier function and reducing inflammation (51). Therefore, the juxtaposition of butyrate and TMA production from γbb highlights the complexity of microbial functions in the context of the human host and the need to fully understand the chemistry underlying metabolite production in order to evaluate the potential impact.

Anaerobic γbb metabolism is a microbial function found in only a select group of human gut bacteria. Prior to this work, *E. timonensis* was the only organism known to perform this transformation and, correspondingly, was found in high abundance in high-TMAO producers from a human carnitine-challenge study (53). However, *E. timonensis* was only detected in 25% of individuals from that group (53), suggesting that additional gut bacteria possess this metabolic activity. Indeed, we have discovered that the *bbu* gene cluster is present in other related gut bacteria. The *bbu* gene cluster was found exclusively in obligate anaerobic bacteria of the Clostridiales order, but it is not ubiquitous within this taxonomic group. This observation contrasts with choline metabolism, which is widely distributed across different gut microbial phyla and classes (54). In addition, the *bbu* gene cluster is only found in host-associated bacteria. While most *bbu*-encoding bacteria were isolated from human stool, a few bacteria originated from the human oral cavity. Bacteria that inhabit the gastrointestinal tract of other animals (*i*.*e*., cat and fish) were also found to possess *bbu* gene clusters, suggesting a broader range of hosts. Anaerobic TMA production from γbb has not been reported outside of host-associated contexts, consistent with our finding that the *bbu* gene cluster is not found in environmental bacteria. Thus, a select group of anaerobic gut bacteria appear to possess a specialized metabolism that supports their growth in this niche.

Our discovery of the genetic basis for γbb metabolism confirms the proposal that the anaerobic conversion of L-carnitine to TMA is an interspecies metabolic pathway. Our analyses show that bacteria possessing the *bbu* pathway to produce TMA from γbb do not have the *cai* operon required to convert L-carnitine to γbb, and vice versa. This observation could explain why γbb-metabolizers inhabit host environments where L-carnitine is abundant due to dietary consumption and where other bacteria can convert it to γbb. The inability of γbb-metabolizing bacteria to generate γbb from L-carnitine highlights the importance of cross-feeding in the gut microbiota. There are now many examples of microbial co-metabolism in the gut (55), including polysaccharide utilization (56), lignin degradation (57), and even drug metabolism (58). Since TMA can only be produced anaerobically when both L-carnitine and γbb utilizing organisms are present, an investigation into their co-occurrence will be important to deconvolute the contribution of anaerobic L-carnitine metabolism to TMA generation in the human gut.

Our initial bioinformatic analyses indicate that the *bbu* pathway producing TMA is likely functionally important in the human gut. Using the *bbuA* gene as a marker for metabolism, we find that this activity is widely distributed in human stool metagenomes. We also detect *bbuA* transcripts in human stool sequencing data, demonstrating that these genes are expressed in the gut environment. Furthermore, the presence of the *bbuA* gene and transcripts is correlated with lower levels of γbb in stool, suggestive of active γbb consumption in subjects possessing this gut bacterial pathway. Conversely, the *cntA* gene is not correlated with changes in stool metabolite levels, nor is it highly expressed in the human gut. This lack of expression could perhaps indicate dioxygen-dependent regulation of *cntA*. Intriguingly, 79% of the HMP2 samples (30/38) that were positive for *cntA* transcripts were from donors with IBD, suggesting that factors associated with inflammation, such as increased O_2_ levels, may induce expression of *cntA*. An independent study also concluded that the *cntA* gene is not expressed in healthy human gut microbiomes (40). Our analysis replicates their reported results using two of the same cohorts (MLVS and iHMP) and adds the larger HMP2 cohort that was since published. These results suggest that the O_2_-independent *bbu* pathway, rather than the O_2_-dependent pathway, is more relevant for TMA production from L-carnitine in the anoxic gut environment. This conclusion warrants caution when inferring functional relevance from gene presence alone and highlights the importance of considering how environmental factors can influence microbial metabolic functions.

Our evaluation of anaerobic γbb metabolism in the human gut microbiota suggests that this pathway may be associated with disease states. In IBD cohorts, the lower prevalence of the *bbuA* gene and transcripts could reflect a decrease in O_2_-intolerant bacteria, such as *bbu*-encoding Clostridiales, due to elevated O_2_ levels caused by inflammation. This trend has been noted for other microbial functions that are restricted to obligate anaerobes, for example metabolism of cholesterol to coprostanol by uncultured *Clostridia* (59). Although TMA production has not been linked to IBD pathology, this observation demonstrates how environmental conditions can influence metabolic functions. On the other hand, the strong connection between TMA production from L-carnitine and CVD motivated our targeted analysis of γbb metabolism in this disease. We did not find a significant difference in the prevalence of *bbu* genes in CVD donors compared to healthy controls, but we did observe that the abundance of *bbu* and *cai* genes was elevated in CVD donors. However, our analysis of paired metagenomic and metatranscriptomic datasets (not associated with CVD) highlights challenges in drawing conclusions from metagenomic data alone. We found that the presence or absence of the *bbu* genes in metagenomic data does not always reflect the presence or absence of *bbu*-encoding organisms. For example, samples that are negative for *bbu* genes can be positive for *bbu* transcripts. Furthermore, we conclusively demonstrate that expression of the *bbu* gene cluster is regulated by γbb and L-carnitine in *E. timonensis*. These results together indicate that transcriptomics is a better reporter of metabolic function, particularly for low-abundance members of the microbiota like γbb-metabolizing bacteria.

A more comprehensive analysis of the contribution of anaerobic γbb metabolism to TMA production in human populations should integrate dietary information and host serum metabolomics with meta-omics analyses. Dietary studies have demonstrated a greater capacity for the gut microbiota from omnivorous donors to produce TMA from L-carnitine compared to vegans or vegetarians (7, 13, 24, 53). Our analysis of the *bbuA* gene in self-reported dietary groups did not reveal differences in prevalence or abundance. However, a more thorough analysis of co-occurrence of L-carnitine and γbb metabolism in controlled dietary cohorts may reveal differences. Diet may also be a complicating factor to consider in meta-analyses of disease cohorts. Finally, although our results show that the *bbuA* gene and transcripts are correlated with lower γbb levels, implying increased production of TMA from this substrate, we were unable to evaluate TMA and TMAO levels from stool metabolomics in these human populations because they are most reliably detected in serum, plasma, or urine. Demonstrating this correlation will be important to conclusively link anaerobic metabolism of L-carnitine-derived γbb to elevated systemic levels of TMA(O) in healthy and disease populations. Our discoveries set the groundwork for subsequent analyses to address this connection.

This work highlights the importance of enzyme discovery for understanding the chemical basis of gut microbial metabolite production, connecting metabolic activities with specific microbes, and analyzing gut microbial functions in human populations. We have uncovered a new enzymatic strategy to generate TMA in an O_2_-independent manner. The BbuA enzyme acts in a pathway that leads to the production of metabolites that are independently associated with human disease and health. In addition, we have discovered a new group of TMA-producing bacteria in the human gut microbiota. The identification of the *bbu* gene cluster in many uncultured bacteria highlights this portion of the microbiota as a rich source of new chemistry that can have important roles in human health. Finally, the discovery of the *bbu* pathway fills an important gap in our knowledge of TMA production from L-carnitine in the anoxic human gut. With a more complete understanding of the metabolic network involved in TMA production, we can begin to dissect the dietary, microbial, and metabolic factors that lead to its generation and connection to human disease.

## ACKNOWLEDGEMENTS

The authors acknowledge funding from the Bill and Melinda Gates Foundation (HHMI-Gates Faculty Scholar Award to E.P.B), a Merck Helen Hay Whitney Foundation fellowship to L.J.R., and the NSF-GRFP fellowship (DGE1144152) to B.F. The authors thank the Broad Microbial ‘Omics Core for RNA isolation, sequencing, and data analysis; Harvard Research Computing for computational resources, maintenance, and support; Dr. Ben Woolston (Northeastern University) for the gift of the pET-proD vector; and Dr. Alain Perret (Institut de biologie François Jacob) for the gift of the *Sinorhizobium meliloti bcoAB* plasmid.

## MATERIALS AND METHODS

### Cloning and plasmids

Genomic DNA (gDNA) was extracted from *E. timonensis* SN18 cultures using the DNeasy UltraClean Microbial Kit (Qiagen, Germantown, MD) according to the manufacturer’s protocol. Oligonucleotide primers were synthesized by Sigma-Aldrich. Genes from the *bbu* gene cluster were amplified from 10 ng *E. timonensis* SN18 gDNA in a 50 μL reaction containing 0.5 μM of forward and reverse primers (**Dataset S5**) and Phusion High-Fidelity PCR Master Mix with HF buffer (New England Biolabs, Ipswich, MA). A plasmid based on the pET28a(+) vector for constitutive expression was constructed using the following protocol and verified by Sanger sequencing. The empty vector pET28a(+) was digested with the restriction enzymes FspI and BamHI (New England Biolabs, Ipswich, MA). After PCR cleanup using the Zymo DNA Clean and Concentrator Kit (Zymo, Irvine, CA), the linearized vector was assembled with a GBlock (**Dataset S5**) carrying the proD promoter (60) using HiFi DNA Assembly Master Mix (New England Biolabs, Ipswich, MA). Vectors used to construct plasmids containing the *bbu* genes were linearized by PCR amplification from 10 ng vector template DNA in a 50 μL reaction containing 0.5 μM of forward and reverse primers (**Dataset S5**) and Phusion High-Fidelity PCR Master Mix with HF buffer (New England Biolabs, Ipswich, MA). Thermocycling was carried out in a Bio-Rad C100 thermal cycler using the following parameters: denaturation at 95 °C for 2 min, followed by 35 cycles of 95 °C for 30 s, 60 °C for 30 s, 72 °C for 1 min/kb, and a final extension at 72 °C for 5 min. PCR reactions were analyzed by agarose gel electrophoresis with ethidium bromide staining and purified using the Illustra GFX PCR DNA and Gel Band Purification kit (GE Healthcare, Chicago, IL). Reactions (10 μL) containing 50 ng linearized vector, 4 molar equivalents of gene insert, and Gibson Assembly Master Mix (New England Biolabs, Ipswich, MA) were incubated at 50 °C for 1 h and transformed into *E. coli* TOP10 chemically competent cells (Thermo Fischer Scientific, Waltham, MA). Transformants were screened by antibiotic selection. Plasmids were purified using the E.Z.N.A. Plasmid Mini Kit I (Omega Bio-tek, Norcross, GA) according to the manufacturer’s protocol and confirmed by Sanger sequencing by Eton Bioscience (Charlestown, MA).

### Bacterial strains and culture conditions

*E. timonensis* SN18 was purchased from Leibniz Institute DSMZ. *E. timonensis* SN18 culturing was performed in Hungate or Balch tubes (Chemglass Life Sciences, Vineland, NJ) at 37 °C and set up in an anaerobic chamber (Coy Laboratory Products, Grasslake, MI) under an atmosphere of 2–4% H_2_, 2–4% CO_2_, and N_2_ as the balance. Standard cultures were grown in PYG medium, modified (medium recipe DSM 104) that was sparged with N_2_ after autoclaving. Basal medium lacking electron acceptors was prepared as described previously (58), containing 1 g/L tryptone (trypticase peptone; BD Biosciences, San Jose, CA), 1 g/L yeast extract (BD Biosciences, San Jose, CA), 0.4 mM L-cysteine, 2.5 g/L NaHCO_3_, 1 g/L NaCl, 0.5 g/L MgCl_2_•6H_2_O, 0.2 g/L KH_2_PO_4_, 0.3 g/L NH_4_Cl, 0.3 g/L KCl, 0.015 g/L CaCl_2_•2H_2_O, 0.25 mL/L of 0.1% resazurin, and 1% ATCC vitamins and trace mineral solutions (American Type Culture Collection, Manassas, VA). NCE medium lacking carbon sources was prepared as described previously (61), containing 4 g/L KH_2_PO_4_, 5 g/L K_2_HPO_4_, 3.5 g/L NaNH_4_PO_4_ 40 mM sodium fumarate dibasic, 1 mM MgSO_4_•7H_2_O, 0.1% casamino acids (VWR Life Science, Radnor, PA), and 1% ATCC vitamin and trace mineral solutions (American Type Culture Collection, Manassas, VA). All chemicals were purchased from Sigma-Aldrich (St. Louis, MO) unless otherwise indicated.

### Chemical synthesis of [*N*-(CD_3_)_3_]-γbb and D_6_-γbb

The following general procedure was used for synthesis of both [*N*-(CD_3_)_3_]-γbb and D_6_-γbb, based on a published protocol (62). In an oven-dried round bottom flask under nitrogen, 1 molar equivalent of GABA (1 g, 9.7 mmol) or D_6_-GABA (92 mg, 0.8 mmol) and 4.6 molar equivalents of K_2_CO_3_ were dissolved in anhydrous methanol (0.1 M) and stirred at room temperature for 15 min. Next, 5.1 molar equivalents of CD_3_I (3.1 mL, 49.4 mmol) or CH_3_I (0.42 mL, 4.3 mmol) was added to the solution, which was stirred at room temperature for 3 days. The reaction mixture was then concentrated under vacuum. The residue was resuspended in chloroform. The solid was filtered under vacuum and washed twice with chloroform. The solid was dissolved in 10% aqueous HCl (0.5 M). The resulting solution was concentrated under vacuum and further dried under vacuum. The residue was triturated three times with acetone. The combined organic layers were concentrated under vacuum. Anhydrous THF (0.1 M) was added and the solution was cooled to 0 °C. The solid was filtered under vacuum and washed with fresh THF, to afford the desired products.

[*N*-(CD_3_)_3_]-γbb (white solid; 160 mg, 1 mmol, **10%**): ^1^H NMR (400 MHz, D_2_O) *δ* ppm 2.07-2.16 (m, 2H), 2.53 (q, 2H, *J* = 7.3 Hz), 3.37 (td, 2H, *J* = 8.3, 3.9 Hz); ^13^C NMR (100 MHz, D_2_O) *δ* Ppm 75.2, 87.3, 109.8, 122.4, 232.4. The NMR data were in accordance with literature values (62).

D_6_-γbb (yellow solid; 76 mg, 0.4 mmol, **48%**): ^1^H NMR (400 MHz, D_2_O) *δ* ppm 3.07 (s, 9H).

### Assays in *E. timonensis* whole cell suspensions

*E. timonensis* SN18 was cultured anaerobically at 37 °C in 10 mL PYG-modified medium supplemented with 1 mM γbb, DL-carnitine, or GABA or with an equivalent volume of 1× PBS. When the cultures reached an OD_600_ = 0.5, they were centrifuged at 1500 × g for 10 min at 4 °C. In an anerobic chamber (Coy Laboratory Products, Grasslake, MI) under an atmosphere of 2-4% H_2_, 2-4% CO_2_, and N_2_ as the balance, the cell pellets were resuspended in 1 mL anoxic 1× PBS and centrifuged at 1500 × g for 10 min at 4 °C. The cell pellets were resuspended in 0.5 mL anoxic 1× PBS and incubated at room temperature for 1 h with 1 mM γbb, [*N*-(CD_3_)_3_]-γbb, D_6_-γbb, DL-carnitine, or GABA or with an equivalent volume of 1× PBS. Reactions were analyzed by LC–MS as described below using three different methods for γbb/TMA, CoA, and fatty acid detection.

### RNA sample preparation, sequencing, and data analysis

*E. timonensis* SN18 cultures were prepared in an anerobic chamber (Coy Laboratory Products, Grasslake, MI), under an atmosphere of 2-4% H_2_, 2-4% CO_2_, and N_2_ as the balance, in Balch tubes containing 15 mL PYG-modified medium. Cultures were grown anaerobically at 37 °C to an OD_600_ = 0.7, when 0.15 mL of a 1 M O_2_-free stock solution of γbb prepared in 1× PBS was added (1 mM final concentration) or an equivalent volume of O_2_-free 1× PBS was added to triplicate cultures for each condition. In a separate experiment (Figure S2), when cultures reached an OD_600_ = 0.5, 0.15 mL of 1 M O_2_-free stock solutions of γbb, DL-carnitine, or GABA prepared in 1× PBS were added (1 mM final concentration) or an equivalent volume of O_2_-free 1× PBS was added to triplicate cultures for each condition. Cultures were grown anaerobically at 37 °C for an additional 45 min and then centrifuged at 3,320 × g for 10 min at 4 °C. Keeping the samples cold on ice, the supernatant was decanted, then removed entirely using a micropipette. Cell pellets were immediately resuspended in 0.5 mL cold Trizol reagent (ThermoFisher, Waltham, MA) and flash frozen in liquid N_2_. Samples were stored at –80 °C until further processing.

Total RNA was isolated by bead beating using 0.1 mm dia. Zirconia/Silica Beads (BioSpec Products, Bartlesville, OK) at 10 m/s for 90 s to lyse cells and then using the Zymo Research Direct-Zol RNA MiniPrep Plus kit (Zymo, Irvine, CA) according to the manufacturer’s protocol. Illumina cDNA libraries were generated using a modified version of the RNAtag-Seq protocol (63). Briefly, 500 ng of total RNA was fragmented, depleted of genomic DNA, and dephosphorylated prior to its ligation to DNA adapters carrying 5′-AN8-3′ barcodes with a 5′ phosphate and a 3′ blocking group. Barcoded RNAs were pooled and depleted of rRNA using the RiboZero rRNA depletion kit (Epicentre, Madison, WI). These pools of barcoded RNAs were converted to Illumina cDNA libraries in three main steps: (i) reverse transcription of the RNA using a primer designed to the constant region of the barcoded adaptor; (ii) addition of a second adapter on the 3’ end of the cDNA during reverse transcription using SmartScribe RT (Clontech Biotechnologies, Mountain View, CA) as previously described (63); (iii) PCR amplification using primers that target the constant regions of the 3’ and 5’ ligated adaptors and contain the full sequence of the Illumina sequencing adaptors. cDNA libraries were sequenced on Illumina HiSeq 2500.

For the analysis of RNAtag-Seq data, reads from each sample in the pool were identified based on their associated barcode using custom scripts, and up to one mismatch in the barcode was allowed with the caveat that it did not enable assignment to more than one barcode. Barcode sequences were removed from the first read as were terminal G’s from the second read that may have been added by SMARTScribe during template switching. Reads were aligned to the *E. timonensis* SN18 genome (NCBI GenBank: FLKM00000000.1) using BWA (64) and read counts were assigned to genes and other genomic features using custom scripts. Differential expression analysis was conducted with DESeq2 (65).

### Heterologous overexpression in *E. coli* and protein purification

Plasmids (**Dataset S5**) containing the *E. timonensis bbuB* gene or the *bbuC* gene in a pET28a(+) vector or the *Sinorhizobium meloloti bcoAB* gene in a pET22 vector (66) were used to transform *E. coli* BL21 Codon-Plus (DE3-*pLys*) chemically competent cells (Invitrogen, Carlsbad, CA). Transformed cells with kanamycin and chloramphenicol resistance were cultured at 37 °C with shaking (180 rpm) in rich Luria Broth (LB) medium with 50 mg/L kanamycin and 25 mg/L chloramphenicol. When cultures reached an OD_600_ between 0.6-0.8, protein expression was induced by addition of IPTG to a final concentration of 0.25 mM. The cultures were then incubated at 15 °C with shaking (180 rpm) for ∼18 h and harvested by centrifugation at 6,000 × *g* for 15 min at 4 °C. Cell pellets were flash frozen in liquid N_2_ and stored at –80 °C until further use. Cells were resuspended in lysis buffer [50 mM *tris*-(hydroxymethyl)aminomethane (Tris)-HCl (pH 7.5) buffer, 150 mM NaCl, and 10% glycerol] at a ratio of approximately 10 mL buffer per 1 g of wet cell mass. The suspension was lysed by passaging twice through a cell disrupter at 12,000 psi and centrifuged at 20,000 × *g* for 30 min at 4 °C. The supernatant was loaded onto a Ni^2+^-NTA immobilized affinity chromatography column (∼10 mL resin per 100 mL lysate) pre-equilibrated with lysis buffer. After loading, 3 column volumes of lysis buffer were passed through the column, followed by 5 column volumes of wash buffer (lysis buffer with 50 mM imidazole). Protein elution was achieved by passing elution buffer (lysis buffer with 250 mM imidazole) over the column. Fractions containing the protein of interest were combined and concentrated at 3,500 × *g* using a 10K MWCO Amicon Ultra-15 Centrifugal Filter Unit (Millipore Sigma, Burlington, MA). The concentrated protein was then dialyzed three times for 4 h each against 100 equivalent volumes of lysis buffer. The protein was frozen in liquid N_2_ and stored at –80 °C. Protein purity was assessed by SDS-PAGE with Coomassie staining, and protein concentration was determined by using the molar absorption coefficient at 280 nm.

*E. coli* BL21 competent cells (Invitrogen, Carlsbad, CA) were co-transformed with a pET28a plasmid containing the *E. timonensis bbuA* gene and a pACYC-Duet1 plasmid containing the *E. timonensis groEL-ES* genes (**Dataset S5**). Transformed cells with kanamycin and chloramphenicol resistance were cultured at 37 °C with shaking (180 rpm) in LB medium with 50 mg/L kanamycin, 25 mg/L chloramphenicol, and 0.2 mM riboflavin. When cultures reached an OD_600_ of 0.6, protein expression was induced by addition of IPTG to a final concentration of 0.25 mM. The cultures were then incubated at 15 °C with shaking (180 rpm) for ∼18 h and harvested by centrifugation at 6,000 × *g* for 15 min at 4 °C. Cell pellets were flash frozen in liquid N_2_ and stored at –80 °C until further use. Cells were resuspended in lysis buffer [50 mM potassium phosphate (pH 7.5) buffer, 300 mM NaCl, 10% glycerol, and 1 mM FAD] at a ratio of approximately 10 mL buffer per 1 g of wet cell mass. The suspension was lysed by passaging twice through a cell disrupter at 12,000 psi and centrifuged at 20,000 × *g* for 30 min at 4 °C. The supernatant was loaded onto a Ni^2+^-NTA immobilized affinity chromatography column (∼10 mL resin per 100 mL lysate) pre-equilibrated with lysis buffer without FAD. After loading the lysate, 3 column volumes of lysis buffer without FAD were passed through the column, followed by 3 column volumes of wash buffer A [50 mM potassium phosphate (pH 7.5) buffer, 300 mM NaCl, 10% glycerol, and 50 mM imidazole] and 3 column volumes of wash buffer B [50 mM potassium phosphate (pH 7.5) buffer, 300 mM NaCl, 10% glycerol, and 75 mM imidazole]. Protein elution was achieved by passing elution buffer [50 mM potassium phosphate (pH 7.5) buffer, 300 mM NaCl, 10% glycerol, and 250 mM imidazole] over the column. Fractions containing the protein of interest were combined and immediately concentrated at 3,500 × *g* using a 30K MWCO Amicon Ultra-15 Centrifugal Filter Unit (Millipore Sigma, Burlington, MA). The protein was frozen in liquid N_2_ without buffer exchange and stored at –80 °C. Protein purity was assessed by SDS-PAGE with Coomassie staining, and protein concentration was determined by using the molar absorption coefficient at 280 nm. FAD-bound protein concentration was determined by using a molar absorption coefficient of 11,300 M^-1^cm^-1^ at 450 nm.

### Chemical synthesis of crotonyl-CoA

Crotonyl-CoA was synthesized according to a published protocol (67) with the following modifications. One molar equivalent of solid Coenzyme A trilithium salt (CoAla Biosciences; 35 mg, 0.046 mmol) was dissolved in 3 mL of ice-cold water. To this solution, 1.5 molar equivalents of crotonic anhydride (Sigma Aldrich; 10 μL, 0.068 mmol) was added and the pH was adjusted to pH 7.0 using a saturated solution of NaHCO_3_. The reaction was stirred in an ice bath for 1.5 h. Reaction progress was monitored using the DTNB “Ellman’s” assay to detect free CoA. The reaction was quenched by adding 48 μL formic acid to lower the pH below 3 and was kept frozen at –20 °C prior to purification. The crotonyl-CoA product was purified using an Thermo Scientific Dionex UltiMate 3000 HPLC instrument and a Thermo Scientific Hypersil GOLD aQ C18 preparative column (250 × 20 mm, 5 μM). The reaction solution was injected onto the column equilibrated with 100% solvent A (10 mM ammonium acetate) and the following gradient method was applied over 21 min with a flow rate of 8 mL/min: 1) solvent B was increased to 40% from 0 to 14 min, 2) solvent B was increased to 95% from 14 to 15 min, 3) solvent B was maintained at 95% from 15 to 16 min, 4) solvent B was decreased to 0% from 16 to 16.5 min, and 5) the column was re-equilibrated with 100% solvent A from 16 to 21 min for subsequent injections. Fractions with absorption at 260 nm were collected and lyophilized to afford crotonyl-CoA as a white solid (10 mg, 0.012 mmol, **26%**). ^1^H NMR (400 MHz, D_2_O) *δ* ppm 0.78 (s, 3H), 0.91 (s 3H), 1.87 (d, 3H, *J* = 6.7 Hz), 2.44 (t, 2H, *J* = 6.1 Hz), 3.03 (t, 2H, *J* = 6.1 Hz), 3.35 (t, 2H, *J* = 6.2 Hz), 3.46 (t, 2H, *J* = 6.2 Hz), 3.58 (dd, 1H, *J* = 4.3 Hz, *J* = 9.3 Hz), 3.85 (dd, 1H, *J* = 4.3 Hz, *J* = 9.6 Hz), 4.04 (s, 1H), 4.26 (br s, 2H), 4.61 (br s, 1H), 4.84-4.87 (m, 2H), 6.17-621 (m, 2H), 6.95 (sextet, 1H, *J* = 7.2 Hz), 8.27 (s, 1H), 8.56 (s, 1H).

### Enzymatic synthesis of γbb-CoA

A 1 mL solution of 0.02 mM *S. meliloti* BcoAB, 0.2 M MgCl_2_, 0.2 M ATP, 0.02 M CoA, 0.05 M γbb in 0.5 M Tris-HCl (pH 8.0) was incubated at room temperature for 30 min. The reaction was quenched with 1% acetic acid (final concentration) and incubated at –20 °C for 30 min. The solution was centrifuged at 16,000 × *g* for 30 min. The γbb-CoA product was purified by preparative HPLC using the same method as described above for crotonyl-CoA. Fractions with absorption at 260 nm were collected and lyophilized. The product identity was confirmed by MS.

### CoA transferase assays in *E. coli* lysates

*E. coli* MG1655 chemically competent cells were transformed with pET-proD plasmids containing *E. timonensis bbu* genes (**Dataset S5**). Transformed cells with kanamycin resistance were grown in 50 mL LB medium with 50 mg/L kanamycin at 37 °C with shaking (180 rpm) for 20 h, reaching an OD_600_ of 1-1.4. Culture aliquots of 15 mL were harvested by centrifugation at 3,220 × g for 15 min at 4 °C and cell pellets were frozen at –80 °C until further use. Cell pellets were resuspended with a volume of 50 mM potassium phosphate (pH 7.5) buffer, 300 mM NaCl, 10% glycerol to achieve a normalized OD_600_ of 20. The cells were lysed on ice by sonication for a total of 1 min, with cycles of 2.5 s on and 10 s off. Reactions (0.05 mL total) containing 0.01 mL of crude lysate, 10 mM acetyl-CoA, 25 mM γbb in 100 mM Tris-HCl (pH 8.0) buffer were incubated at room temperature for 1 h and analyzed by LC–MS using the CoA detection method.

### TMA-lyase assays in *E. coli* lysates

*E. coli* MG1655 (DE3) chemically competent cells were transformed with a pET28a plasmid containing *E. timonensis bbu* genes and the pACYC-Duet1 plasmid containing the *E. timonensis groEL-ES* genes (**Dataset S5**). Transformed cells with kanamycin and chloramphenicol resistance were grown in 50 mL LB medium with 50 mg/L kanamycin, 25 mg/L chloramphenicol, and 0.2 mM riboflavin at 37 °C with shaking (180 rpm). When cultures reached an OD_600_ of 0.6, protein expression was induced by addition of IPTG to a final concentration of 0.25 mM. The cultures were then incubated at 15 °C with shaking (180 rpm) for ∼15 h. Cultures aliquots of 15 mL were harvested by centrifugation at 3,220 × g for 15 min at 4 °C and cell pellets were frozen at –80 °C until further use. Cell pellets were transferred and thawed in an anerobic chamber (Coy Laboratory Products; Grasslake, MI) located in a cold room at 4 °C under an atmosphere of ∼3% H_2_ and N_2_ as the balance. The pellets were resuspended with a volume of O_2_-free 50 mM potassium phosphate (pH 7.5) buffer, 300 mM NaCl, 10% glycerol to achieve an OD_600_ = 10. The cells were lysed by sonication for a total of 1 min, with cycles of 2.5 s on and 10 s off. In an anaerobic chamber (Coy Laboratory Products; Grasslake, MI) under an atmosphere of ∼3% H_2_ and N_2_ as the balance, reactions (0.05 mL) containing 0.02 mL of crude lysate, 10 mM acetyl-CoA, 10 mM γbb in O_2_-free 100 mM Tris-HCl (pH 8.0) buffer were incubated at room temperature for 1 h and analyzed by LC–MS using the methods for CoA and TMA detection.

### CoA transferase *in vitro* activity assay

Reactions (50 μL) containing 0.01 mM BbuB, 0.01 mM BbuC, 10 mM acetyl-CoA, and 10 mM γbb in 100 mM Tris-HCl (pH 8.0) buffer were incubated at room temperature for 1 h and analyzed by LC–MS using the CoA detection method.

### TMA-lyase *in vitro* activity assay

Protein solutions of BbuA, BbuB, and BbuC were deoxygenated on ice by eight rapid cycles of vacuum and N_2_ gas for a total of four times. In an anaerobic chamber (Coy Laboratory Products; Grasslake, MI) under an atmosphere of ∼3% H_2_ and N_2_ as the balance, reactions (0.1 mL) containing 0.2 mM BbuA, 0.01 mM BbuB, 0.01 mM BbuC, 10 mM acetyl-CoA, and 10 mM γbb in O_2_-free 100 mM Tris-HCl (pH 8.0) buffer were incubated at room temperature for 1 h and analyzed by LC–MS using the methods for CoA and TMA detection.

### Liquid chromatography – mass spectrometry sample preparation and analytical methods

*γbb and TMA analysis*. Samples were diluted 1:10 (*v/v*) in solvent A composed of 95:5 acetonitrile (ACN):100 mM ammonium formate (MS grade) in H_2_O with 0.02% formic acid (MS grade) and centrifuged at 3,220 × *g* for 10 min. The extracted supernatant samples were further diluted 1:4 (*v/v*) in solvent A and kept at 4 °C until analysis (performed within 24 h).

Liquid chromatography was conducted using an Agilent 1200 Series G1312B Binary Pump (Agilent Technologies, Santa Clara, CA) instrument and a Phenomenex HILIC column (30 mm × 2.1 mm, 2.5 μm). Samples (2 μL) were injected onto the column equilibrated with 100% solvent A and the following gradient method was applied over 7.5 min with a flow rate of 0.6 mL/min: 1) Solvent A was maintained at 100% from 0 to 1 min, 2) solvent B was increased to 90% from 1 to 3 min, 3) solvent B was maintained at 90% from 3 to 4.5 min, 4) solvent B was decreased to 0% from 4.5 to 6 min, and 5) the column was re-equilibrated with 100% solvent A from 6 to 7.5 min. Tandem MS/MS detection was performed with an Agilent 6410 Triple Quadrupole (Agilent Technologies, Santa Clara, CA) instrument with electron spray ionization in positive mode (ESI+). The source parameters were gas temperature of 200 °C, gas flow of 10 L/min, nebulizer pressure of 45 psi, and capillary voltage of 4000 V. Conditions for MS/MS fragmentation for the metabolites of interest are found in **Table 1**. Standards were prepared according to the same methods used for experimental samples and standard curves were used to quantify metabolite concentrations.

**Table 1.** MS/MS parameters used to detect metabolites of interest.

| Metabolite | Parent ion<br>[M+H]<sup>+</sup> (*m/z*) | Daughter ion<br>[M+H]<sup>+</sup> (*m/z*) | Dwell (ms) | Fragmentor (V) | CE (V) |
| --- | --- | --- | --- | --- | --- |
| **γbb** | 146 | 87 | 200 | 110 | 21 |
| **TMA** | 60 | 45 | 200 | 110 | 26 |
| **DL-Carnitine** | 162 | 103 | 200 | 110 | 21 |
| **GABA** | 104 | 87 | 200 | 110 | 21 |

#### Derivatization of fatty acids with 2-nitrophenylhydrazine/1-ethyl-3-(3-dimethylaminopropyl)-carbodiimide and analysis

Samples were diluted 1:10 (*v/v*) in 95:5 ACN:100 mM ammonium formate (MS grade) in H_2_O with 0.02% formic acid (MS grade) and centrifuged at 3,220 × *g* for 10 min. The extracted supernatant samples (10 μL) were diluted 1:10 (*v/v*) in a solution of 10% 1:1 pyridine:HCl (pH 3.5-5) in H_2_O. A pre-mixed derivatization solution consisting of equal parts 0.29 M 1-ethyl-3-(3-dimethylaminopropyl)-carbodiimide (EDC) hydrochloride (dissolved in water) and 0.12 M 2-nitrophenylhydrazine (2-NPH; dissolved in 0.25 M HCl) was prepared fresh and 20 μL of this solution was added to 100 μL of 100-fold diluted sample. The derivatization reactions were heated at 60 °C for >15 min and then diluted 1:10 (*v/v*) in H_2_O prior to LC–MS analysis.

Liquid chromatography was conducted using an Agilent 1200 Series G1312B Binary Pump (Agilent Technologies, Santa Clara, CA) instrument and an Agilent Extend C18 column (50 mm × 2.1 mm, 1.8 μm). The derivatized samples (2 μL) were injected onto the column equilibrated with 95% H_2_O (solvent A) and 5% ACN (solvent B). The following gradient method was applied over 13 min with a flow rate of 0.3 mL/min: 1) solvent A was maintained at 95% from 0 to 4 min, 2) solvent B was increased to 100% from 0 to 4 min, 3) solvent B was maintained at 100% from 4 to 7 min, 4) solvent B was decreased to 5% from 7 to 9 min, and 5) the column was re-equilibrated with 95% solvent A from 9 to 13 min. An MS scan was performed with an Agilent 6410 Triple Quadrupole (Agilent Technologies, Santa Clara, CA) instrument with electron spray ionization in negative mode (ESI–). The source parameters were a gas temperature of 325 °C, a gas flow of 12 L/min, a nebulizer pressure of 40 psi, and a capillary voltage of 4000 V. The detection mode was set to a scan range of 100–400 *m/z* and a scan time of 500 ms with a fragmentor voltage of 135 V and a step size of 0.1 amu. The parent [M-H]^−^ ions for derivatized short-chain fatty acids were monitored, including formate (180 *m/z*), acetate (194 *m/z*), propionate (208 *m/z*), and butyrate (222 *m/z*). Butyrate concentrations were determined by comparison to a 10 μM sodium ^13^C_4_-butyrate internal standard added to the derivatization solution.

#### CoA-thioester analysis

Cell cultures (5 mL) or whole cell suspensions (1.5 mL) were centrifuged at 3,220 × *g* or 16,000 × *g*, respectively, for 10 min at 4 °C. Cell pellets were resuspended in 0.2 mL of cold solution of 40:20:20 ACN:MeOH:H_2_O. Resuspended cells were incubated at –20 °C for 30 min and then centrifuged at 16,000 × *g* for 30 min. The supernatant was transferred to a new tube and dried using a Genevac EZ-2.3 Elite Evaporation System (Genevac Ltd, Ipswich, UK) for 1 h. The dried samples were resuspended in 80 μL H_2_O for LC–MS analysis. Samples (20 μL) from lysate or *in vitro* reactions were quenched in 2 μL concentrated acetic acid, diluted 1:10 in H_2_O, and incubated at –20 °C for 30 min before centrifugation at 16,000 × *g* for 10 min. Samples were further diluted 1:4 in H_2_O and kept at 4 °C prior to LC–MS analysis. Standards were prepared according to the same methods used for experimental samples.

Liquid chromatography was conducted using a Waters Acquity UPLC H-Class System (Waters Corporation, Milford, MA) instrument and a Waters Acquity UPLC BEH C18 column (2.1 × 50 mm, 1.7 μm). The samples (2 μL) were injected onto the column equilibrated with 100% 30 mM ammonium acetate (solvent A). The following gradient method was applied over 5 min with a flow rate of 0.8 mL/min: 1) solvent A was maintained at 100% from 0 to 0.5 min, 2) solvent B (acetonitrile + 0.1 % formic acid) was increased to 20% from 0.5 to 4 min, 3) solvent B was increased to 100% from 4.0 to 4.1 min, 4) solvent B was maintained at 100% from 4.1 to 4.4 min, 5) solvent B was decreased to 0% from 4.4 to 4.5 min, and 5) the column was re-equilibrated with 100% solvent A from 4.5 to 5 min. MS detection was performed with a Waters Xevo TQ-S (Waters Corporation, Milford, MA) instrument with electron spray ionization in positive mode (ESI+). The source parameters were gas temperature of 500 °C, desolvation gas flow of 1000 L/h, cone gas flow of 150 L/h nebulizer pressure of 7 bar, capillary voltage of 0.5 kV and cone voltage of 3 V. Conditions for tandem MS/MS detection were optimized using an acetyl-CoA standard. Conditions for a parent MS/MS scan of the 136 *m/z* mass fragment (corresponding to adenine loss) were a scan range of 800–910 *m/z*, scan time of 0.15 s, collision energy of 53 V, and cone voltage of 70 V. Conditions for an MS scan were a scan range of 800–910 *m/z*, cone voltage of 70 V, and scan time of 0.15 s. Conditions for MS/MS fragmentation for the metabolites of interest are found in **Table 2**.

**Table 2.** MS/MS parameters used to detect metabolites of interest.

| Metabolite | Parent ion<br>[M+H]<sup>+</sup> (*m/z*) | Daughter ion<br>[M+H]<sup>+</sup> (*m/z*) | Dwell (ms) | Fragmentor (V) | CE (V) |
| --- | --- | --- | --- | --- | --- |
| Acetyl-CoA | 810.1957 | 135.9617 | 52 | 70 | 54 |
| Crotonyl-CoA | 836.5 | 135.9617 | 52 | 70 | 54 |
| Butyryl-CoA | 838.12 | 135.9617 | 52 | 70 | 54 |
| 3-Hydroxybutyryl-CoA | 854.0 | 135.9617 | 52 | 70 | 54 |
| γbb-CoA | 895.5 | 135.9617 | 52 | 70 | 54 |

### Bioinformatic analyses

#### Genome Searches

The *E. timonensis* BbuA protein sequence was used in a Basic Local Alignment Search Tool (BLAST) search of the National Center for Biotechnology Information (NCBI) non-redundant protein database, the UniProt database (release 2019_07), and the Joint Genome Institute-Integrated Microbial Genomes (JGI-IMG) database of all isolates. Hits with >70% amino acid sequence identity were considered BbuA homologs. Searches were also conducted of the HMP1 reference genomes and genomes from the human gut bacteria culture collections described in **Dataset S3**. No additional hits were identified from these searches. Next, collections of human metagenome-assembled genomes (MAGs) (**Dataset S3**) were performed using a tblastn search and an e-value cutoff of <0.0001. Hits were manually evaluated and further filtered using a >70% amino acid sequence identity cutoff. Contigs containing hits were annotated using the Galaxy webtool Prokka (68, 69).

#### Phylogenetics

A multiple sequence alignment was generated using MAFT v7.455 (70) of the BbuA homolog protein sequences and 2,388 representatives of protein clusters with >80% amino acid sequence ID from the top 10,000 hits of a BLAST search of the UniProt database (release 2019_07) using the *E. timonensis* SN18 BbuA protein as a query. A maximum-likelihood phylogenetic tree was constructed using IQ-TREE v1.6.12 (71) with the LG+F0+G12 model and visualized using FigTree v1.4.4 (72). Branch supports were calculated using the aBayes method (73).

#### Metagenome and Metatranscriptome Searches & Quantification

BbuA homolog protein sequences (**Dataset S6**) were used to generate a protein database. A negative control protein database was made using protein sequences acquired from the follow steps: (1) the top 10,000 results were obtained from a BLAST search of the UniProt database (release 2019_07) using *E. timonensis* SN18 BbuA as the query, excluding BbuA homologs described above, (2) these sequences were clustered using UniRef50 (release 2019_07), resulting in 288 representative proteins, and (3) each unique sequence from step 1 between 50-70% sequence identity to *E. timonensis* SN18 BbuA was added to the list from step 2. The CntA protein database consisted of *E. coli* YeaW (UniProt P0ABR8), *A. baumannii* CntA (UniProt D0C9N6), and *Klebsiella pneumoniae* CntA (UniProt A0A377WGT7).

Human studies reporting the stool meta’omic data that were analyzed in this work are listed in **Dataset S4**. A blastx DIAMOND(74) search with an e-value cut-off of <0.0001 and a percent amino acid sequence identity of >50% was performed using the raw shotgun metagenome or metatranscriptome sequencing reads against the BbuA, CntA, and negative control databases. If the highest sequence identity hit for the read to a BbuA protein was greater than or equal to that of a negative control protein <u>and</u> the sequence identity to the BbuA protein was >70%, then that read was considered a positive hit for a *bbuA* gene or transcript. The positive hits for each metagenome sample were summed and then normalized by RPKM and average genome size (AGS) using the following equations.

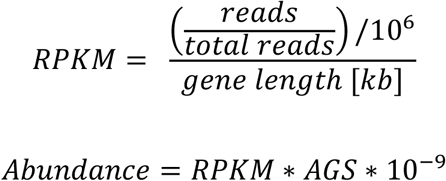

Average genome size for each sample was calculated using MicrobeCensus (75). Plots and statistical analyses were performed using the ggplot2 package v3.3.2 (76) and R v3.6.0 (77).

## SUPPORTING INFORMATION

The Supporting Information includes additional figures (S1-S15) and legends for datasets (S1-S6) that are provided as separate files.

## Notes

### Competing Interest Statement

The authors have declared no competing interest.

## REFERENCES

1. E. Holmes, et al., Understanding the role of gut microbiome-host metabolic signal disruption in health and disease. Trends Microbiol 19, 349–359 (2011).

2. J. K. Nicholson, et al., Host-gut microbiota metabolic interactions. Science 336, 1262–1267 (2012).

3. M. S. Donia & M. A. Fischbach, Small molecules from the human microbiota. Science 349, 1254766 (2015).

4. M. al-Waiz, M. Mikov, S. C. Mitchell, R. L. Smith, The exogenous origin of trimethylamine in the mouse. Metabolism 41, 135–136 (1992).

5. C. J. Rebouche & H. Seim, Carnitine metabolism and its regulation in microorganisms and mammals. Annu Rev Nutr 18, 39–61 (1998).

6. Z. Wang, et al., Gut flora metabolism of phosphatidylcholine promotes cardiovascular disease. Nature 472, 57–63 (2011).

7. R. A. Koeth, et al., Intestinal microbiota metabolism of L-carnitine, a nutrient in red meat, promotes atherosclerosis. Nat Med 19, 576–585 (2013).

8. L. Hoyles, et al., Metabolic retroconversion of trimethylamine N-oxide and the gut microbiota. Microbiome 6, 73 (2018).

9. Z. Wang, et al., Prognostic value of choline and betaine depends on intestinal microbiota-generated metabolite trimethylamine-N-oxide. Eur Heart J 35, 904–910 (2014).

10. D. H. Lang, et al., Isoform specificity of trimethylamine N-oxygenation by human flavin-containing monooxygenase (FMO) and P450 enzymes: selective catalysis by FMO3. Biochem. Pharmacol. 56, 1005–1012 (1998).

11. S. C. Mitchell & R. L. Smith, Trimethylaminuria: The Fish Malodor Syndrome. Drug Metab Dispos 29, 517–521 (2001).

12. S. H. Zeisel & M. Warrier, Trimethylamine N-Oxide, the microbiome, and heart and kidney disease. Annu Rev Nutr 37, 157–181 (2017).

13. R. A. Koeth, et al., L-Carnitine in omnivorous diets induces an atherogenic gut microbial pathway in humans. J Clin Invest 129, 373–387 (2019).

14. J. A. Meadows & M. J. Wargo, L-Carnitine in bacterial physiology and metabolism. Microbiology 161, 1161–1174 (2015).

15. C. J. Rebouche, Kinetics, pharmacokinetics, and regulation of L-carnitine and acetyl-L-carnitine metabolism. Ann N Y Acad Sci 1033, 30–41 (2004).

16. A. K. Sawicka, G. Renzi, R. A. Olek, The bright and the dark sides of L-carnitine supplementation: a systematic review. J Int Soc Sports Nutr 17, 49 (2020).

17. B. Wassef, M. Kohansieh, A. N. Makaryus, Effects of energy drinks on the cardiovascular system. World J Cardiol 9, 796–806 (2017).

18. C. J. Rebouche & C. A. Chenard, Metabolic fate of dietary carnitine in human adults: Identification and quantification of urinary and fecal metabolites. J. Nutr. 121, 539–546 (1991).

19. C. J. Rebouche, D. L. Mack, P. F. Edmonson, L-Carnitine dissimilation in the gastrointestinal tract of the rat. Biochemistry 23, 6422–6426 (1984).

20. R. A. Koeth, et al., γ-Butyrobetaine is a proatherogenic intermediate in gut microbial metabolism of L-carnitine to TMAO. Cell Metab 20, 799–812 (2014).

21. Y. Zhu, et al., Carnitine metabolism to trimethylamine by an unusual Rieske-type oxygenase from human microbiota. Proc Natl Acad Sci U S A 111, 4268–4273 (2014).

22. S. Rath, B. Heidrich, D. H. Pieper, M. Vital, Uncovering the trimethylamine-producing bacteria of the human gut microbiota. Microbiome 5, 54 (2017).

23. L. Albenberg, et al., Correlation between intraluminal oxygen gradient and radial partitioning of intestinal microbiota. Gastroenterology 147, 1055–1063 e1058 (2014).

24. W. K. Wu, et al., Identification of TMAO-producer phenotype and host-diet-gut dysbiosis by carnitine challenge test in human and germ-free mice. Gut 68, 1439–1449 (2019).

25. W. Buckel, Enzymatic reactions involving ketyls: From a chemical curiosity to a general biochemical mechanism. Biochemistry 58, 5221–5233 (2019).

26. D. M. Smith, W. Buckel, H. Zipse, Deprotonation of enoxy radicals: theoretical validation of a 50-year-old mechanistic proposal. Angew Chem Int Ed Engl 42, 1867–1870 (2003).

27. S. Craciun & E. P. Balskus, Microbial conversion of choline to trimethylamine requires a glycyl radical enzyme. Proc Natl Acad Sci U S A 109, 21307–21312 (2012).

28. J. R. Andreesen, Glycine reductase mechanism. Curr Opin Chem Biol 8, 454–461 (2004).

29. A. Maurer, F. Leisinger, D. Lim, F. P. Seebeck, Structure and mechanism of ergothionase from Treponema denticola. Chemistry 25, 10298–10303 (2019).

30. S. Ghisla & C. Thorpe, Acyl-CoA dehydrogenases: A mechanistic overview. Eur. J. Biochem. 271, 494–508 (2004).

31. U. Müh, I. Cinkaya, S. P. Albracht, W. Buckel, 4-Hydroxybutyryl-CoA dehydratase from Clostridium aminobutyricum: characterization of FAD and iron-sulfur clusters involved in an overall non-redox reaction. Biochemistry 35, 11710–11718 (1996).

32. H. Seedorf, et al., The genome of Clostridium kluyveri, a strict anaerobe with unique metabolic features. Proc Natl Acad Sci U S A 105, 2128–2133 (2008).

33. J. K. Hardman & T. C. Stadtman, Metabolism of ω-amino acids: I. Fermentation of γ-aminobutyric acid by Clostridium aminobutyricum n. sp. J. Bacteriol. 79, 544–548 (1960).

34. J. Bader, et al., Utilization of (E)-2-butenoate (crotonate) by Clostridium kluyveri and some other Clostridium species. Arch Microbiol 125, 159–165 (1980).

35. A. Almeida, et al., A new genomic blueprint of the human gut microbiota. Nature 568, 499–504 (2019).

36. E. Pasolli, et al., Extensive unexplored human microbiome diversity revealed by over 150,000 genomes from metagenomes spanning age, geography, and lifestyle. Cell 176, 649–662 e620 (2019).

37. J. Lloyd-Price, et al., Multi-omics of the gut microbial ecosystem in inflammatory bowel diseases. Nature 569, 655–662 (2019).

38. M. Schirmer, et al., Dynamics of metatranscription in the inflammatory bowel disease gut microbiome. Nat Microbiol 3, 337–346 (2018).

39. R. S. Mehta, et al., Stability of the human faecal microbiome in a cohort of adult men. Nat Microbiol 3, 347–355 (2018).

40. S. Rath, et al., Pathogenic functions of host microbiota. Microbiome 6, 174 (2018).

41. M. Poyet, et al., A library of human gut bacterial isolates paired with longitudinal multiomics data enables mechanistic microbiome research. Nat Med 25, 1442–1452 (2019).

42. E. A. Franzosa, et al., Gut microbiome structure and metabolic activity in inflammatory bowel disease. Nat Microbiol 4, 293–305 (2019).

43. Z. Jie, et al., The gut microbiome in atherosclerotic cardiovascular disease. Nat Commun 8, 845 (2017).

44. P. Strandwitz, et al., GABA-modulating bacteria of the human gut microbiota. Nat Microbiol 4, 396–403 (2019).

45. D. T. Thwaites, et al., Gamma-aminobutyric acid (GABA) transport across human intestinal epithelial (Caco-2) cell monolayers. Br J Pharmacol 129, 457–464 (2000).

46. U. Scherf, et al., Succinate-ethanol fermentation in Clostridium kluyveri: purification and characterisation of 4-hydroxybutyryl-CoA dehydratase/vinylacetyl-CoA Δ3-Δ2-isomerase. Arch Microbiol 161, 239–245 (1994).

47. H. Huber, et al., A dicarboxylate/4-hydroxybutyrate autotrophic carbon assimilation cycle in the hyperthermophilic Archaeum Ignicoccus hospitalis. Proc Natl Acad Sci U S A 105, 7851–7856 (2008).

48. I. A. Berg, D. Kockelkorn, W. Buckel, G. Fuchs, A 3-hydroxypropionate/4-hydroxybutyrate autotrophic carbon dioxide assimilation pathway in archaea. Science 318, 1782–1786 (2007).

49. J. Zhang, et al., Substrate-induced radical formation in 4-hydroxybutyryl coenzyme A dehydratase from Clostridium aminobutyricum. Appl Environ Microbiol 81, 1071–1084 (2015).

50. S. Bodea, M. A. Funk, E. P. Balskus, C. L. Drennan, Molecular basis of C-N bond cleavage by the glycyl radical enzyme choline trimethylamine-lyase. Cell Chem Biol 23, 1206–1216 (2016).

51. G. R. Nicolas & P. V. Chang, Deciphering the chemical lexicon of host-gut microbiota interactions. Trends Pharmacol Sci 40, 430–445 (2019).

52. G. den Besten, et al., The role of short-chain fatty acids in the interplay between diet, gut microbiota, and host energy metabolism. J Lipid Res 54, 2325–2340 (2013).

53. W. K. Wu, et al., Characterization of TMAO productivity from carnitine challenge facilitates personalized nutrition and microbiome signatures discovery. Microbiome 8, 162 (2020).

54. A. Martinez-del Campo, et al., Characterization and detection of a widely distributed gene cluster that predicts anaerobic choline utilization by human gut bacteria. mBio 6, (2015).

55. C. M. Rath & P. C. Dorrestein, The bacterial chemical repertoire mediates metabolic exchange within gut microbiomes. Curr Opin Microbiol 15, 147–154 (2012).

56. J. M. Grondin, et al., Polysaccharide utilization loci: Fueling microbial communities. J. Bacteriol 199, e00860–00816 (2017).

57. E. N. Bess, et al., Genetic basis for the cooperative bioactivation of plant lignans by Eggerthella lenta and other human gut bacteria. Nat Microbiol 5, 56–66 (2020).

58. V. M. Rekdal, et al., Discovery and inhibition of an interspecies gut bacterial pathway for Levodopa metabolism. Science 364, (2019).

59. D. J. Kenny, et al., Cholesterol metabolism by uncultured human gut bacteria influences host cholesterol level. Cell Host Microbe 28, 245–257 e246 (2020).

60. J. H. Davis, A. J. Rubin, R. T. Sauer, Design, construction and characterization of a set of insulated bacterial promoters. Nucleic Acids Res. 39, 1131–1141 (2011).

61. K. A. Romano, et al., Metabolic, epigenetic, and transgenerational effects of gut bacterial choline consumption. Cell Host Microbe 22, 279–290 e277 (2017).

62. C. Morano, X. Zhang, L. D. Fricker, Multiple Isotopic Labels for Quantitative Mass Spectrometry. Anal. Chem. 80, 9298–9309 (2008).

63. A. A. Shishkin, et al., Simultaneous generation of many RNA-seq libraries in a single reaction. Nat. Meth. 12, 323–325 (2015).

64. H. Li & R. Durbin, Fast and accurate short read alignment with Burrows–Wheeler transform. Bioinformatics 25, 1754–1760 (2009).

65. M. I. Love, W. Huber, S. Anders, Moderated estimation of fold change and dispersion for RNA-seq data with DESeq2. Genome Biol 15, 550 (2014).

66. P. Bazire, et al., Characterization of L-carnitine metabolism in Sinorhizobium meliloti. J. Bacteriol. 201, e00772–00718 (2019).

67. E. J. Simon & D. Shemin, The Preparation of S-Succinyl Coenzyme A. J Am Chem Soc 75, 2520–2520 (1953).

68. T. Seemann, Prokka: rapid prokaryotic genome annotation. Bioinformatics 30, 2068–2069 (2014).

69. G. Cuccuru, et al., Orione, a web-based framework for NGS analysis in microbiology. Bioinformatics 30, 1928–1929 (2014).

70. K. Katoh, K. Misawa, K. Kuma, T. Miyata, MAFFT: a novel method for rapid multiple sequence alignment based on fast Fourier transform. Nucleic acids research 30, 3059–3066 (2002).

71. L.-T. Nguyen, H. A. Schmidt, A. von Haeseler, B. Q. Minh, IQ-TREE: A fast and effective stochastic algorithm for estimating maximum-likelihood phylogenies. Mol Biol Evol 32, 268–274 (2014).

72. A. Rambaut (2012) FigTree v1. 4.

73. M. Anisimova, et al., Survey of Branch Support Methods Demonstrates Accuracy, Power, and Robustness of Fast Likelihood-based Approximation Schemes. Syst Biol 60, 685–699 (2011).

74. B. Buchfink, C. Xie, D. H. Huson, Fast and sensitive protein alignment using DIAMOND. Nat Meth 12, 59–60 (2015).

75. S. Nayfach & K. S. Pollard, Average genome size estimation improves comparative metagenomics and sheds light on the functional ecology of the human microbiome. Genome Biol 16, 51 (2015).

76. H. Wickham (2016) ggplot2: Elegant Graphics for Data Analysis (Springer-Verlag, New York).

77. R. C. Team (2017) R: A language and environment for statistical computing (R Foundation for Statistical Computing, Vienna, Austria).

