## Supporting Information for "Elucidation of an anaerobic pathway for metabolism of L-carnitine-derived γ-butyrobetaine to trimethylamine in human gut bacteria"

**This PDF file includes:**

Figures S1 to S15
Legends for Datasets S1 to S6

**Other supplementary materials for this manuscript include the following:**

Datasets S1 to S6

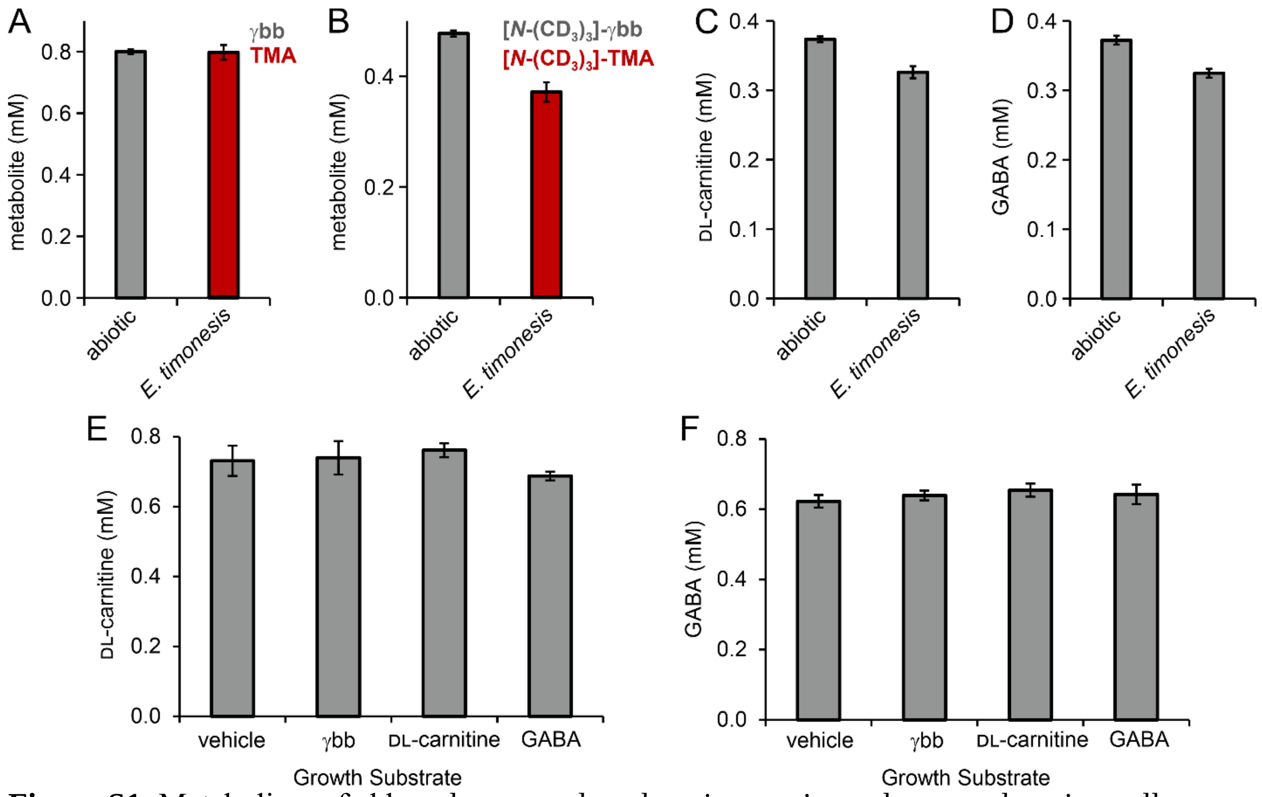


**Figure S1.** Metabolism of γbb and structural analogs in growing cultures and resting cell suspensions of *E. timonensis*. **(A-D)** Stacked bar plots of metabolite concentrations detected by LC-MS from *E. timonensis* SN18 culture extracts after 20 h of growth. Media were supplemented with **(A)** 0.8 mM γbb, **(B)** 0.4 mM [*N*-$(CD_3)_3$]-γbb, **(C)** 0.4 mM DL-carnitine, or **(D)** 0.4 mM GABA. **(E-F)** Concentration of metabolites detected by LC-MS from extracts of resting cell suspensions of *E. timonensis* SN18 that were grown in the presence of 1 mM γbb, DL-carnitine, GABA, or 1× PBS and incubated with **(E)** 0.8 mM DL-carnitine, or **(F)** 0.8 mM GABA. Error bars represent standard deviation from the mean of three biological replicates.

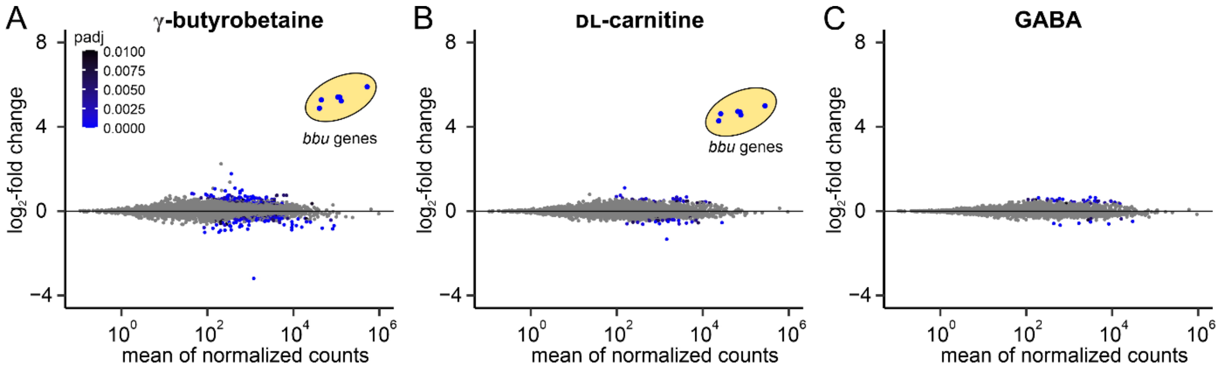


**Figure S2.** Differential gene expression from *E. timonesis* SN18 cultures supplemented with (**A**) γbb, (**B**) DL-carnitine, or (**C**) GABA at an OD = 0.5 compared to cultures treated with a vehicle* plotted against the mean of normalized counts from three biological replicates of the substrate-induced samples. The *bbu* genes are circled in yellow. Genes with an adjusted *p*-value >0.01 (Wald test) comparing γbb- and vehicle-treated cells are represented by grey circles.

*This set of triplicate samples for each condition were conducted in independent experiments from the data presented in Figure 2 of the main text.

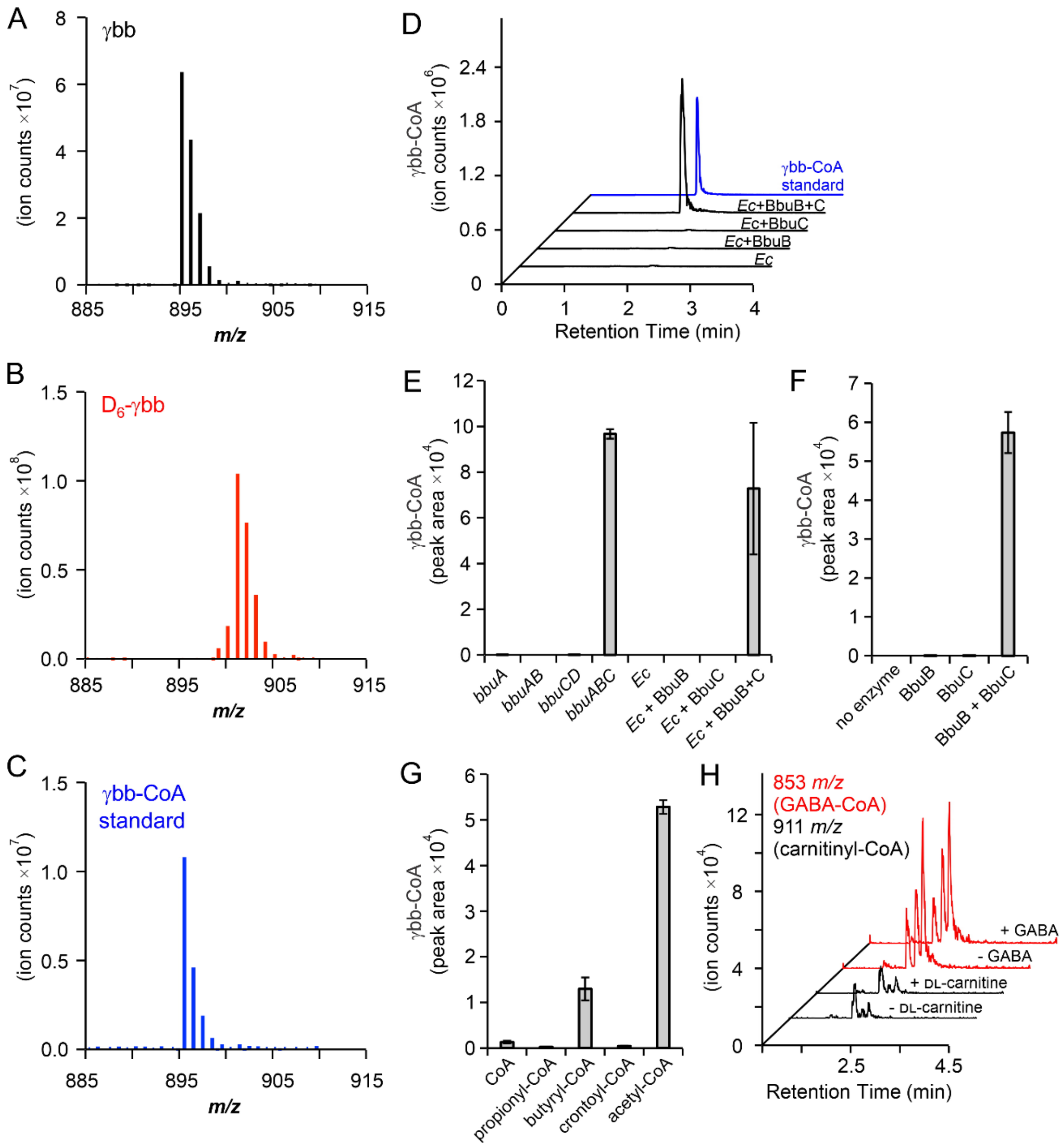


**Figure S3.** Production of γbb-CoA *in vivo* and *in vitro*. Mass spectra of γbb-CoA from extracts of *E. timonensis* cell suspensions incubated for 40 min with (**A**) γbb or (**B**) $D_6$-γbb compared to (**C**) a γbb-CoA standard. (**D**) LC–MS/MS selected ion chromatograms of the 136 *m/z* fragment ion of γbb-CoA from 1 h incubations of γbb, acetyl-CoA, and purified BbuB and BbuC with crude lysate of *E. coli* transformed with an empty vector. Relative amounts of γbb-CoA determined by LC–MS peak area from (**E**) crude lysate and (**F**) *in vitro* assays. (**G**) Relative amounts of γbb-CoA determined by LC–MS peak area using various short-chain fatty acid CoA donor substrates. (**H**) LC–MS/MS selected ion chromatograms of the 136 *m/z* fragment-parent ion pairs for DL-carnitinyl-CoA and GABA-CoA from 3-h incubations of DL-carnitine or GABA and acetyl-CoA with purified BbuB and BbuC. Error bars shown in panels E–G represent standard deviation from the mean of three biological replicates.

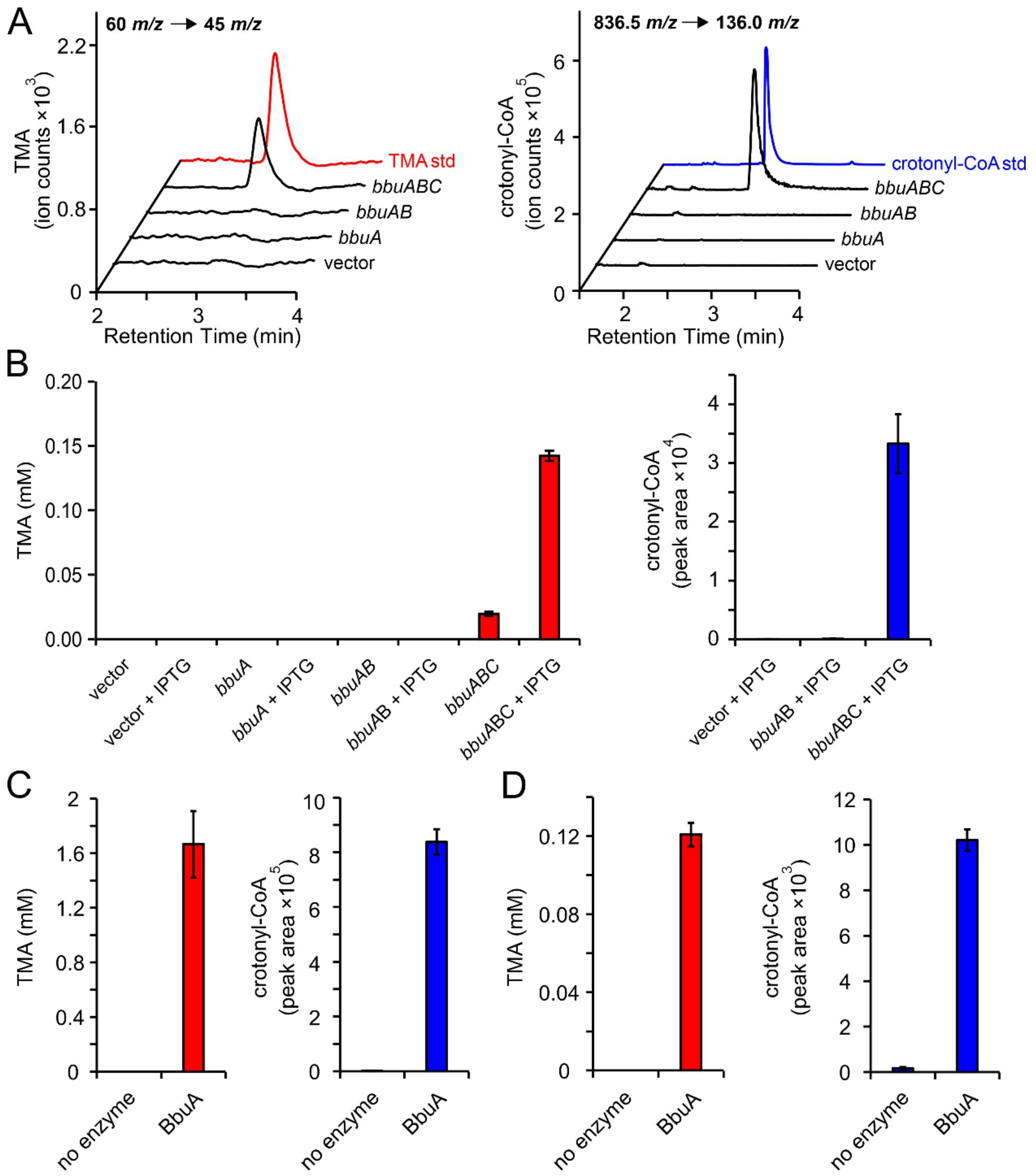


**Figure S4.** Production of TMA and crotonyl-CoA from *E. coli* gain-of-function experiments and *in vitro* activity assays. (**A**) LC–MS/MS selected ion chromatograms (SIC) of the 136.0 *m/z* fragment-precursor ion pairs of TMA and crotonyl-CoA that were produced from 1 h incubations of γbb and acetyl-CoA with crude lysate of IPTG-induced *E. coli* expressing *bbu* genes or empty vector. (**B**) TMA concentrations and crotonyl-CoA relative amounts determined by LC–MS/MS that were produced from 1 h incubations of γbb and acetyl-CoA with crude lysate of IPTG-induced *E. coli* expressing *bbu* genes or empty vector. (**C**) TMA concentrations and crotonyl-CoA relative amounts determined by LC–MS/MS that were produced from 1 h reactions containing γbb, acetyl-CoA, BbuB, BbuC, and FAD with or without addition of BbuA. (**D**) TMA concentrations and crotonyl-CoA relative amounts determined by LC–MS/MS that were produced from 1 h reactions containing γbb-CoA and FAD with or without addition of BbuA.

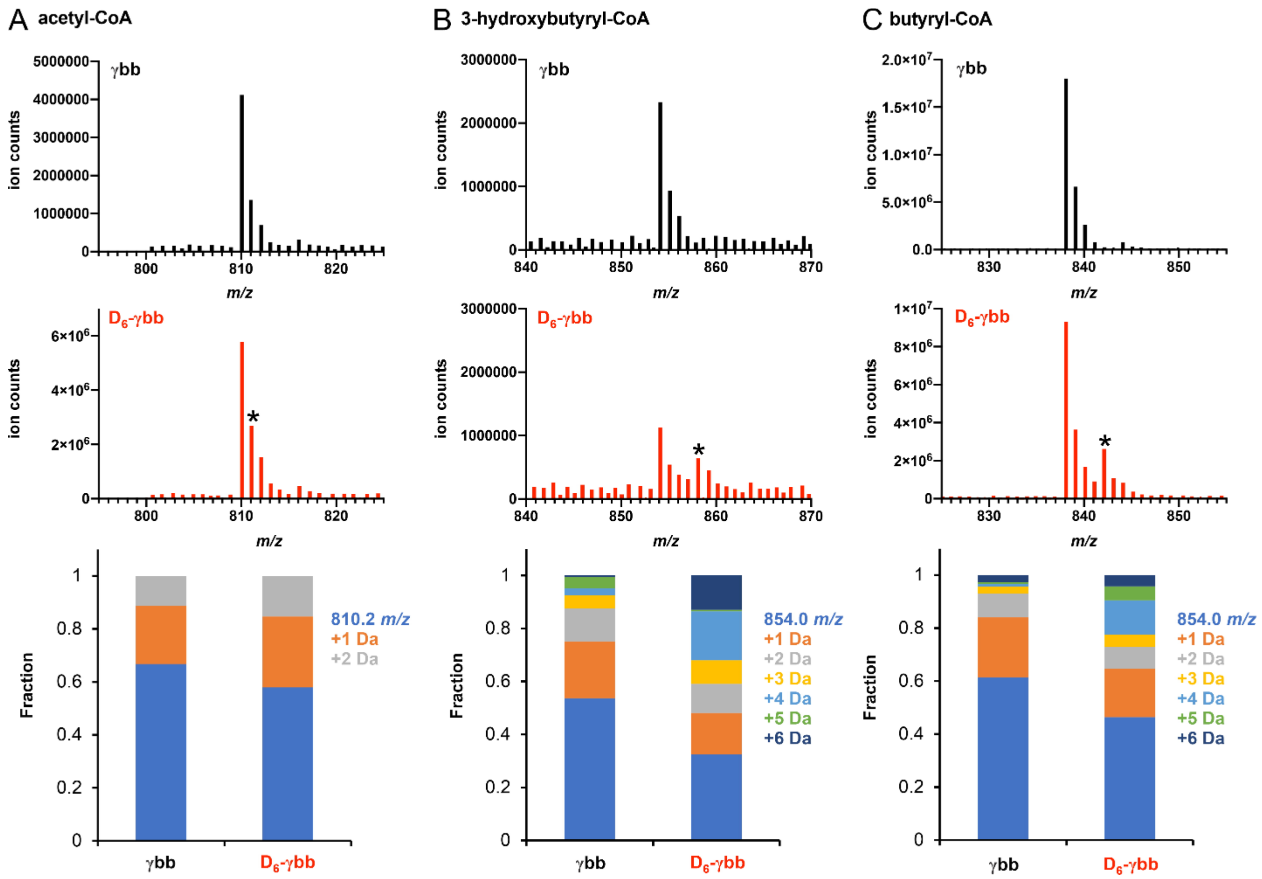


**Figure S5.** Mass spectra of the products of crotonyl-CoA metabolism [(**A**) acetyl-CoA, (**B**) 3-hydroxybutyryl-CoA, (**C**) butyryl-CoA] from extracts of *E. timonensis* SN18 cell suspensions incubated for 40 min with γbb (black) or $D_6$-γbb (red). Asterisks indicate increased abundance of ions attributed to deuterium-labeled isotopologs. Stacked bar plots show fraction of metabolite isotopologs.

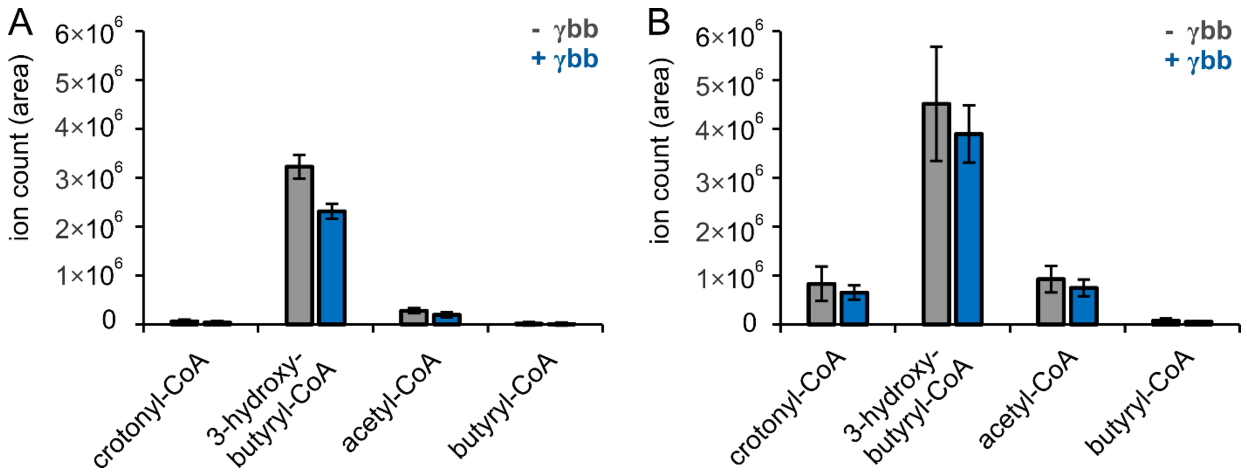


**Figure S6.** Metabolites detected by LC–MS from 1 h (**A**) and 4 h (**B**) incubations of crotonyl-CoA with crude lysate of *E. timonensis* SN18 that was cultured in the presence (blue) or absence (grey) of γbb. Error bars represent the standard deviation from the mean of three biological replicates.

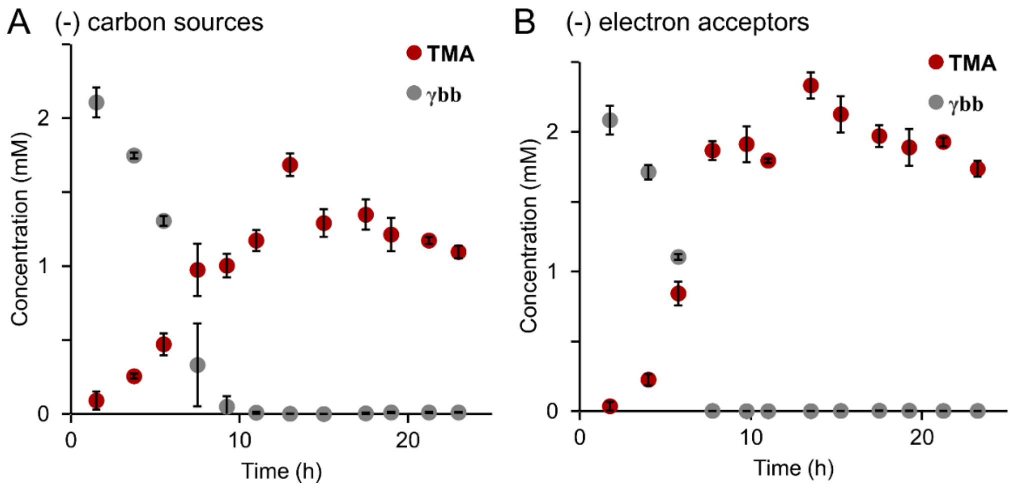


**Figure S7.** Metabolites detected by LC-MS from cell extracts of anaerobic cultures of *E. timonensis* SN18 in media lacking (**A**) carbon sources or (**B**) electron acceptors and containing 2 mM γbb. Error bars represent standard deviation from the mean of three biological replicates.

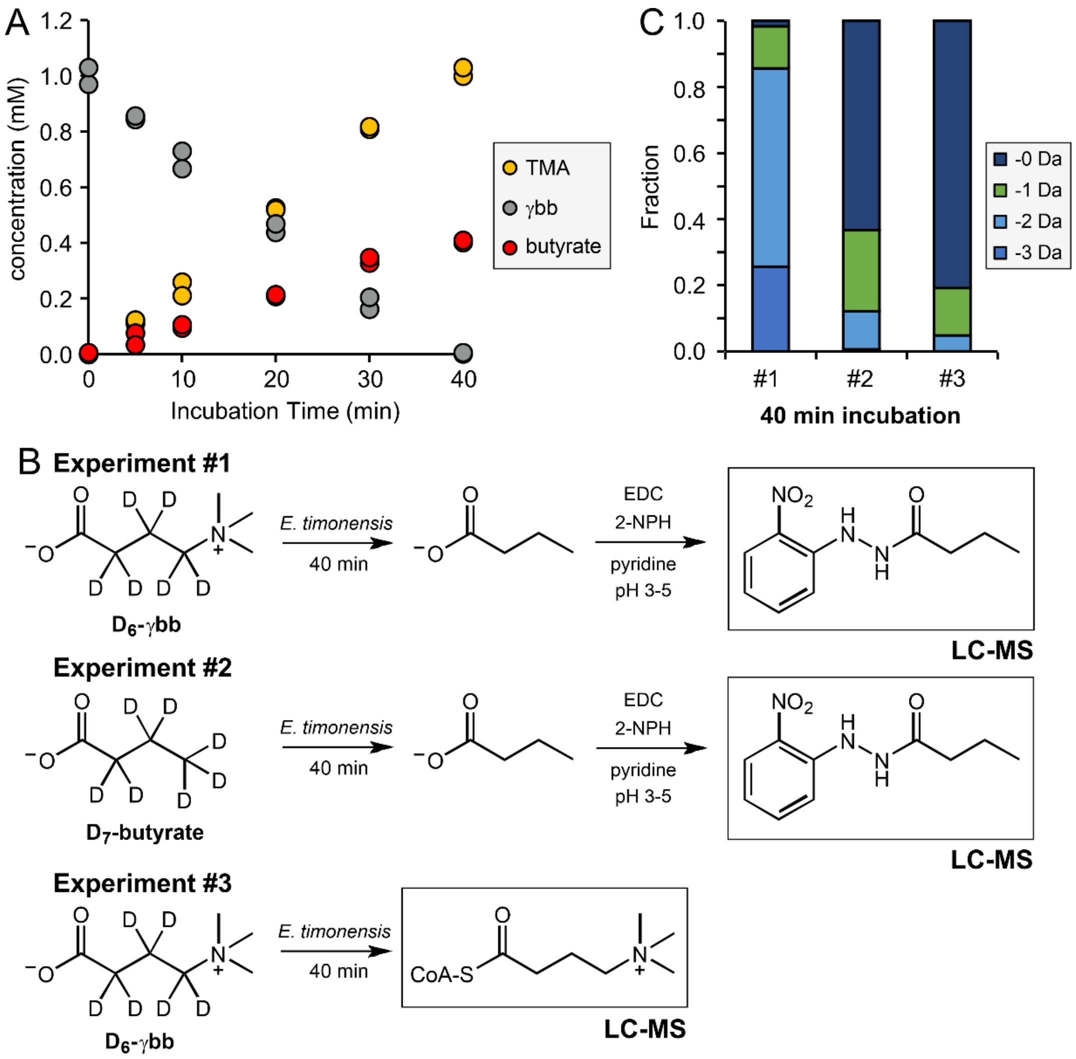


**Figure S8.** (**A**) Butyrate production during an incubation of 1 mM γbb in a resting suspension of *E. timonesis* SN18 cells grown in the presence of γbb. (**B**) Experimental conditions for results shown in panel C. Experiment #1 represents the mass spectrum of derivatized butyrate from a resting cell suspension of γbb-induced *E. timonesis* SN18 incubated with 1 mM $D_6$-γbb for 40 min. Experiment #2 represents the mass spectrum of derivatized butyrate from a resting cell suspension of γbb-induced *E. timonesis* SN18 incubated with 1 mM $D_7$-butyrate for 40 min. Experiment #3 represents the mass spectrum of γbb-CoA from a resting cell suspension of γbb-induced *E. timonesis* SN18 incubated with 1 mM $D_6$-γbb for 40 min. (**C**) Isotopolog distribution of metabolite products detected by LC–MS from the three experiments described in panel B.

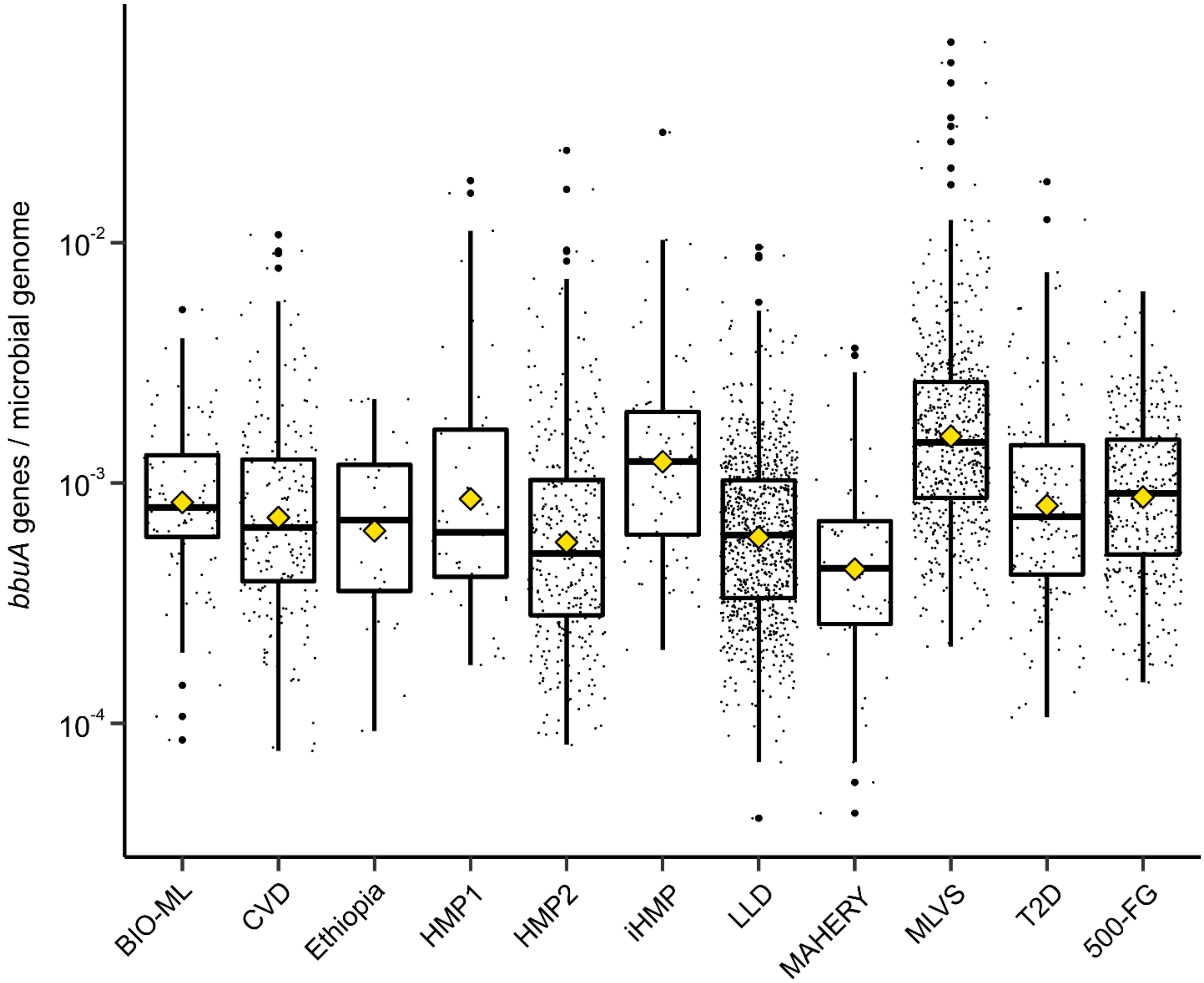


**Figure S9.** Abundance of the *bbuA* gene in stool metagenomes collected from various human studies. Mean values are represented by yellow diamonds.

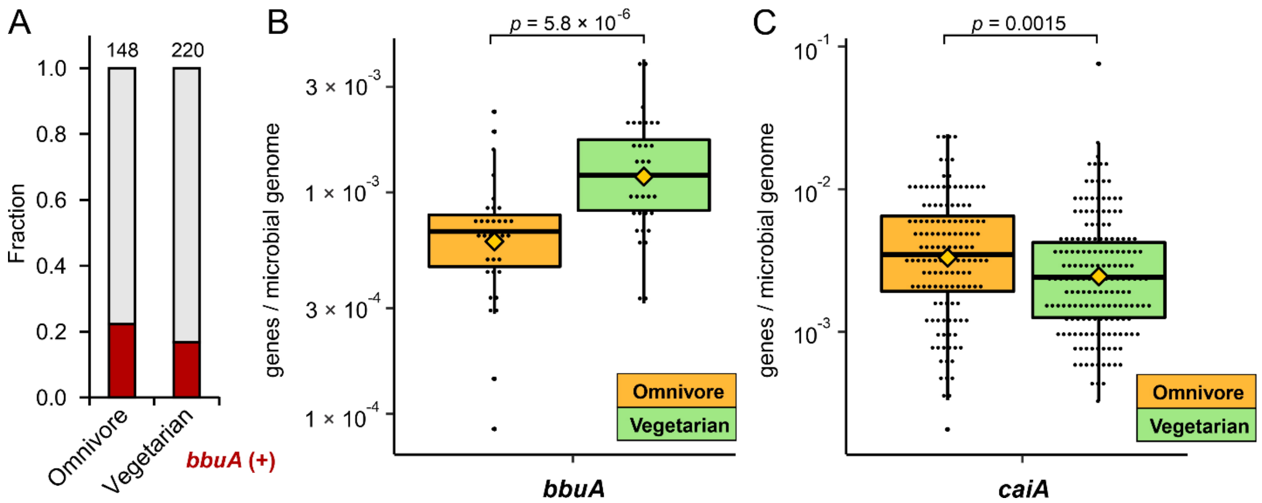


**Figure S10.** Correlation between γbb metabolism and diet. (**A**) Fraction of samples positive and negative for the *bbuA* gene in stool metagenomes of self-reported omnivores and vegetarians from the BIO-ML study. Comparison of (**B**) *bbuA* or (**C**) *caiA* gene abundance in stool metagenomes of self-reported omnivores and vegetarians from the BIO-ML study. Mean values are represented by yellow diamonds and *p*-values were determined using the Mann-Whitney *U*-test.

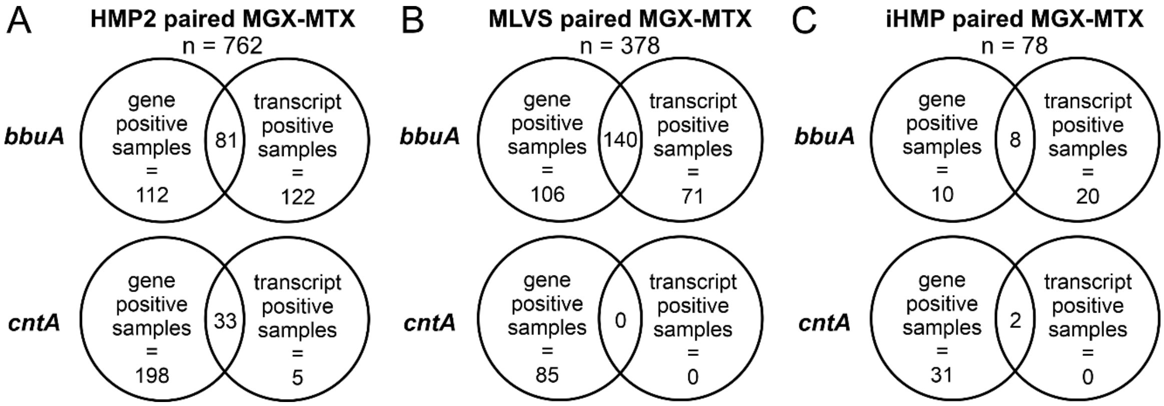


**Figure S11.** Overlap of stool samples from the (**A**) HMP2, (**B**) MLVS, and (**C**) iHMP cohorts that are positive for the *bbuA* or *cntA* gene and/or transcripts.

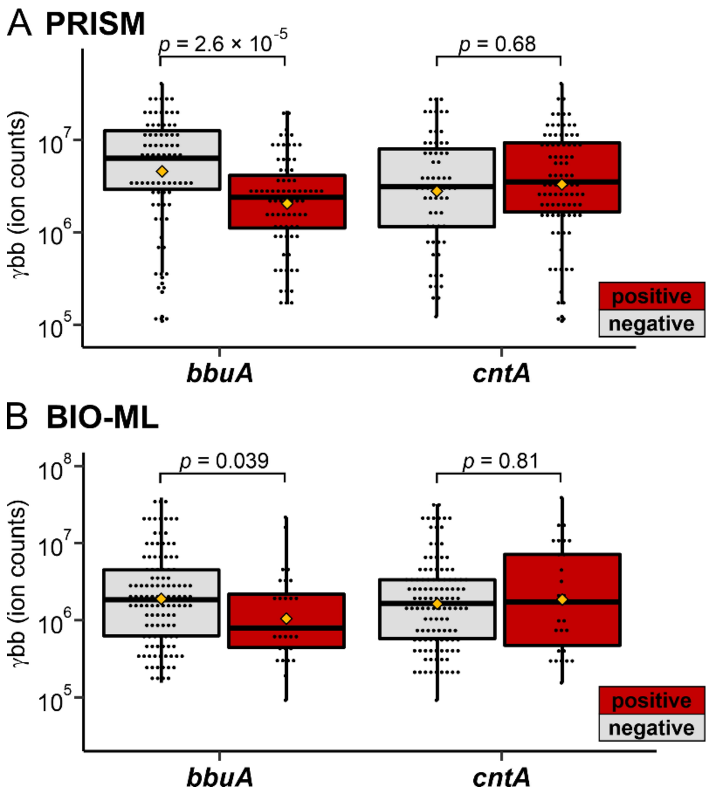


**Figure S12.** Correlations between the presence of *bbuA* or *cntA* in metagenomes and γbb metabolite levels in stool samples from the (**A**) PRISM and (**B**) BIO-ML cohorts. Mean values are represented by yellow diamonds and *p*-values were determined using the Mann-Whitney *U*-test.

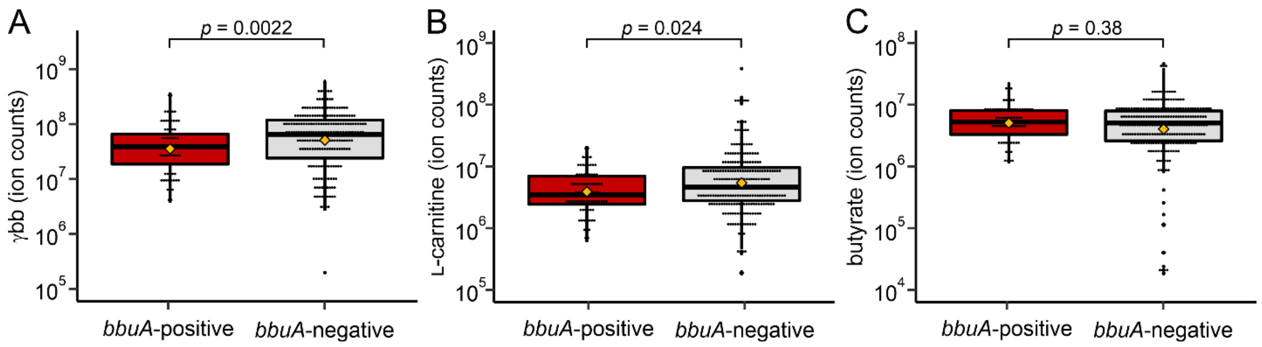


**Figure S13.** Correlations between the presence of *bbuA* in metatranscriptomes and levels of (**A**) γbb, (**B**) L-carnitine and (**C**) butyrate in stool metabolomes from the HMP2 projects. Mean values are represented by yellow diamonds and *p*-values were determined using the Mann-Whitney *U*-test.

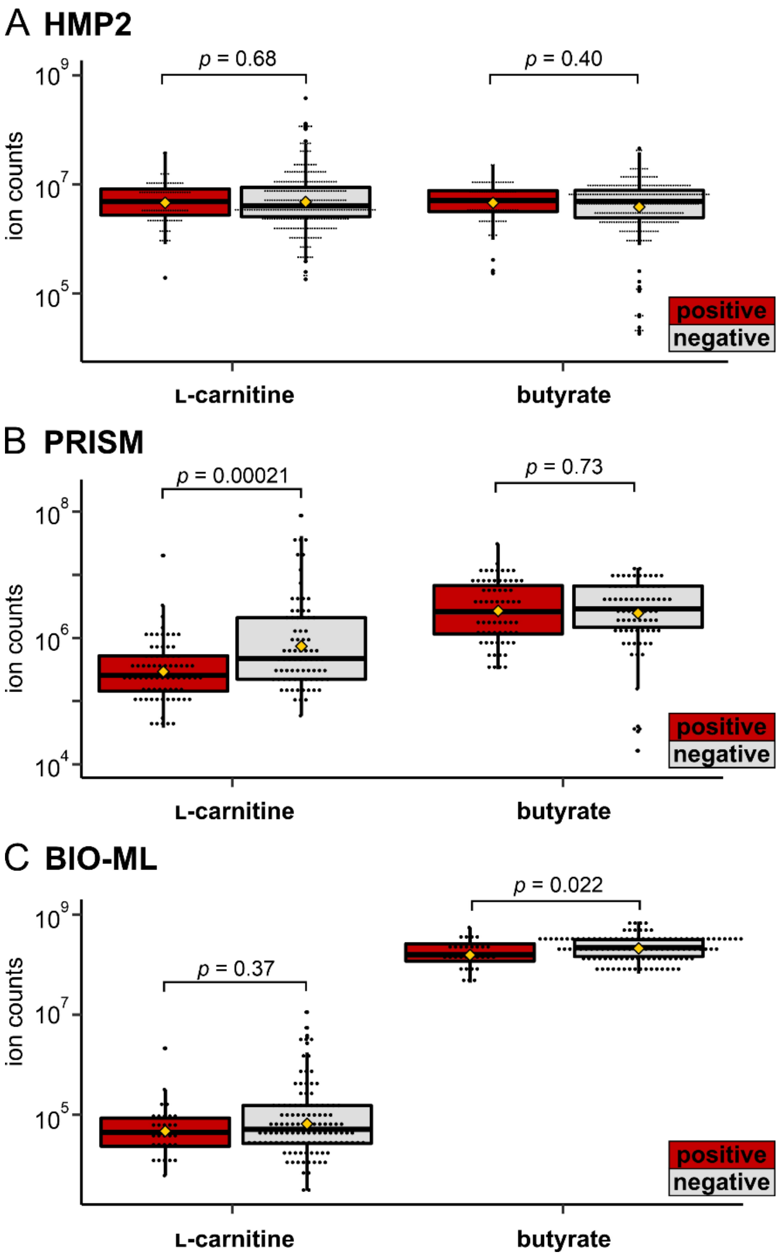


**Figure S14.** Correlations between the presence of *bbuA* in metagenomes and L-carnitine (orange) or butyrate (green) metabolite levels in stool samples from the (**A**) HMP2, (**B**) PRISM, and (**C**) BIO-ML projects. Mean values are represented by yellow diamonds and *p*-values were determined using the Mann-Whitney *U*-test.

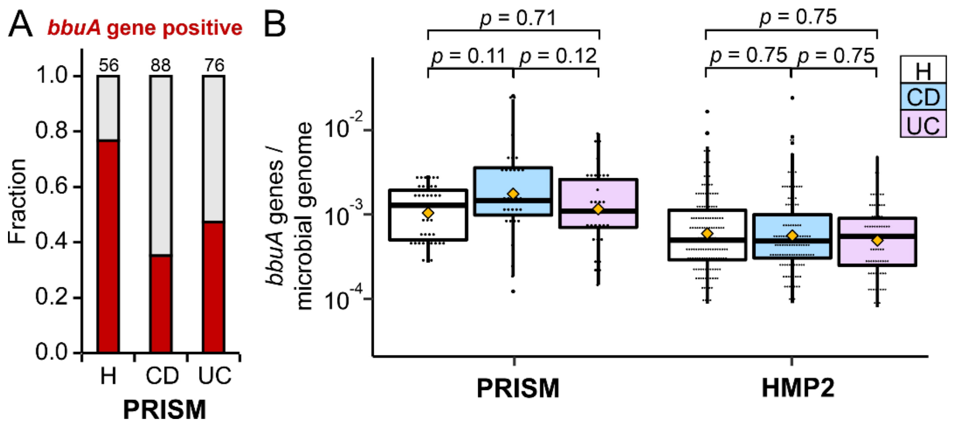


**Figure S15.** (**A**) Proportion of samples positive or negative for the *bbuA* gene in the PRISM Crohn's disease (CD) and ulcerative colitis (UC) cohorts compared to healthy (H) controls. (**B**) Abundance of the *bbuA* gene in IBD cohorts compared to healthy controls in the PRISM and HMP2 studies. Mean values are represented by yellow diamonds and *p*-values were determined using the Mann-Whitney *U*-test.

**Dataset S1 (separate file).** *E. timonensis* RNA-sequencing data statistics and gene expression (RPKM) data.

**Dataset S2 (separate file).** Homologs of enzymes responsible for crotonyl-CoA metabolism in *E. timonensis* and corresponding gene expression data from RNA-seq.

**Dataset S3 (separate file).** List of isolate genome collections and MAGs analyzed for the present of the *bbu* genes. MAGs containing *bbu* gene clusters are provided, including information about study and sample of origin, completeness, contamination, and taxonomy classification.

**Dataset S4 (separate file).** List of human studies with metagenomics datasets used for bioinformatic analyses.

**Dataset S5 (separate file).** Primer sequences and accession codes of gene sequences used in molecular cloning.

**Dataset S6 (separate file).** BbuA protein sequences used for bioinformatic analyses.
